# A broad-host-range CRISPRi toolkit for silencing gene expression in *Burkholderia*

**DOI:** 10.1101/618413

**Authors:** Andrew M. Hogan, A. S. M. Zisanur Rahman, Tasia J. Lightly, Silvia T. Cardona

## Abstract

Genetic tools are critical to dissecting the mechanisms governing cellular processes, from fundamental physiology to pathogenesis. Members of the genus *Burkholderia* have potential for biotechnological applications but can also cause disease in humans with a debilitated immune system. The lack of suitable genetic tools to edit *Burkholderia* GC-rich genomes has hampered the exploration of useful capacities and the understanding of pathogenic features. To address this, we have developed CRISPR interference (CRISPRi) technology for gene silencing in *Burkholderia*, testing it in *B. cenocepacia*, *B. multivorans* and *B. thailandensis*. Tunable expression was provided by placing a codon-optimized *dcas9* from *Streptococcus pyogenes* under control of a rhamnose-inducible promoter. As a proof of concept, the *paaABCDE* operon controlling genes necessary for phenylacetic acid degradation was targeted by plasmid-borne sgRNAs, resulting in near complete inhibition of growth on phenylacetic acid as the sole carbon source. This was supported by reductions in *paaA* mRNA expression. The utility of CRISPRi to probe other functions at the single cell level was demonstrated by knocking down *phbC* and *fliF*, which dramatically reduces polyhydroxybutyrate granule accumulation and motility, respectively. As a hallmark of the mini-CTX system is the broad host-range of integration, we putatively identified 67 genera of Proteobacteria that might be amenable to modification with our CRISPRi toolkit. Our CRISPRi tool kit provides a simple and rapid way to silence gene expression to produce an observable phenotype. Linking genes to functions with CRISPRi will facilitate genome editing with the goal of enhancing biotechnological capabilities while reducing *Burkholderia’*s pathogenic arsenal.

**Author contributions:** STC conceived the idea and design of the research; AMH designed and cloned the dCas9 constructs; AMH and ASMZ designed the sgRNAs, assessed knockdown phenotypes, processed data, and wrote and edited the manuscript; TJL performed RT-qPCR analysis and edited the manuscript; STC supervised the work and provided financial support.

**Graphical Abstract:** 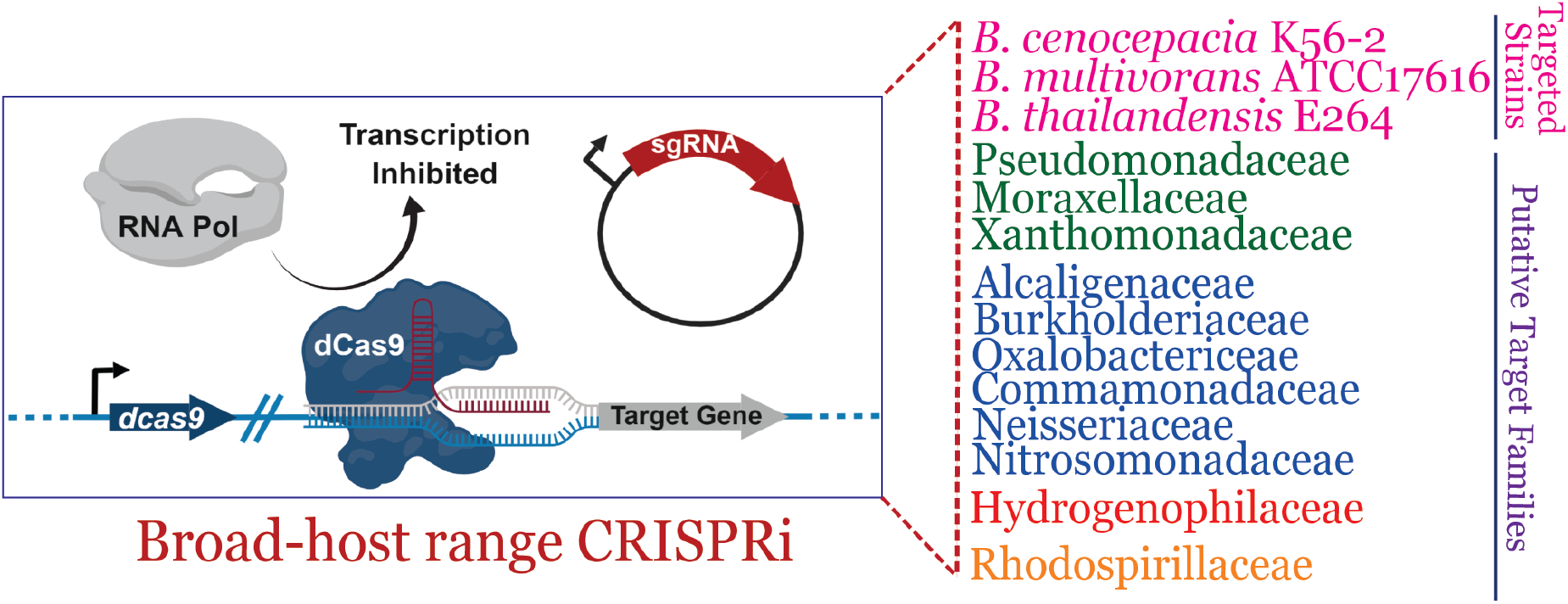

---

The genus *Burkholderia* comprises a diverse group of Gram-negative bacteria characterized by a remarkable biotechnological potential, with species that can be exploited for bioremediation purposes, production of bioactive compounds, and agricultural growth promotion. Perhaps due to the phylogenetic diversity of the genus, some species of *Burkholderia* can also pose a threat to human health ^1^. Within the multiple deep-branching groups of the *Burkholderia* phylogenetic tree ^2^, one branch contains the clades known as the *Burkholderia cepacia* complex (Bcc) and the *B. pseudomallei* group. The Bcc group comprises species that cause severe infections in people with the genetic disease cystic fibrosis and other patients with a compromised immune system ^3^. *B. cenocepacia* and *B. multivorans* are two relevant species of this group, which are prevalent in cystic fibrosis patients ^4^. The *B. pseudomallei* group contains the risk group 3 members *B. mallei*, *B. pseudomallei*, and its close relative *B. thailandensis*, the last one being a frequent surrogate used in *B. pseudomallei* and *B. mallei* research ^5^. Other branches, such as the *B. xenovorans* group comprise species isolated from diverse environmental sources, like polluted soils, and plant rhizospheres ^2^. While species with and without pathogenic potential tend to cluster separately in most phylogenetic trees, the genetic characteristics that define the pathogenic potential of *Burkholderia* are poorly understood. Environmental species can cause serious infections ^6^, calling for caution in the use of *Burkholderia* strains for biotechnological applications. The incomplete understanding of *Burkholderia* pathogenic potential may be related to the limited tools available to link gene to function. Due to high resistance to the antibiotics typically used as genetic markers ^7^, and the high GC content of their large genomes, most genome editing methods designed for Gram-negative bacteria are inefficient and need to be adapted for use in *Burkholderia*.

Tools that facilitate controlled gene expression are necessary for interrogation of gene function. Programmable control of gene expression by promoter replacement is a valuable tool to link gene to phenotype ^8^. Yet, promoter replacement implies that the natural regulatory circuitry of the target gene is interrupted. Instead, clustered regularly interspaced short palindromic repeats interference (CRISPRi) ^9^ is a method of silencing native gene expression, which is based on a dead Cas9 (dCas9) and a nuclease-inactive version of the RNA-guided endonuclease Cas9 ^10^. In CRISPRi, a single guide RNA (sgRNA) designed towards the 5’ end of the target gene and the dCas9 protein form an RNA-protein complex that recognizes the target region by base-pairing, and sterically blocks transcription initiation, if targeting the promoter, or elongation if targeting downstream of the promoter, by the RNA polymerase ^9^. Initially developed in *Escherichia coli*, CRISPRi technology has been adapted to a wide range of bacterial strains to address focused efforts such as metabolic rewiring and antimicrobial characterization ^11–15^, and broader efforts to functionally characterize genomes ^16–18^. To achieve fine control of gene expression, a wide range of dCas9 expression levels are necessary, which can be provided by expressing *dcas9* with strong inducible promoters ^18^ or by providing multiple copies of *dcas9* engineered in a plasmid ^16^. Limitations to the success of these efforts are the proteotoxicity of dCas9 when expressed at high levels ^19^ and the necessity of customizing CRISPRi delivery tools across bacteria. Recently, the use of Tn*7* transposon mutagenesis, which specifically delivers genetic constructs close to a single *glmS* site ^20,21^ was applied to deliver a mobile CRISPRi system across bacteria ^22^. However, the Tn*7* system is less suitable for *Burkholderia* species as their genomes contain multiple copies of *glmS*, requiring additional steps to confirm the site of chromosomal integration ^23^.

In this work, we employ a mini-CTX-derived mutagenesis system ^24^ to achieve specific chromosomal delivery of *dcas9* in three species of *Burkholderia*, *B. cenocepacia*, *B. thailandensis* and *B. multivorans*. By placing the chromosomal copy of *dcas9* under the control of the *E. coli* rhamnose-inducible promoter ^25^, we demonstrate durable and tunable control of endogenous gene expression in *Burkholderia* species, which affects cellular function producing observable phenotypes. We extend the usability of our CRISPRi tool kit by exploring other bacterial genomes for putative mini-CTX insertion sites.

## Results

### Construction of the CRISPRi system

The dCas9 from *Streptococcus pyogenes* has been shown to provide robust gene repression in diverse bacteria ^9,18,22^; we therefore selected it as our first approach. To function, the dCas9 binds a sgRNA and, by complementary base pairing, is guided to target and silences a gene of interest by sterically blocking the RNA polymerase ^9^. However, the genome of *S. pyogenes* has low GC-content (~40%), and from inspection of the *dcas9* gene, we expected poor codon usage in high GC-content organisms such as *Burkholderia* (67%) and subsequently low levels of expression. We first attempted to express the native gene from *S. pyogenes* in single copy in the chromosome under control of the rhamnose-inducible promoter, which is known to yield robust and tightly-regulated expression in *Burkholderia* ^25^; however, we did not observe detectable levels of dCas9 protein expression by immunoblot (Supplemental Figure 1). Upon codon optimization for *B. cenocepacia*, we first introduced the gene into a multicopy plasmid under the control of the rhamnose-inducible promoter ^25^; however, we observed a severe growth defect upon induction, except at minute concentrations of rhamnose (Supplemental Figure 2A). Growth inhibition was not observed in the vector control (Supplemental Figure 2B) and it remains unclear if the inhibitory effect of dCas9 expression on growth was caused by metabolic load from expression of a large protein in multicopy, or from proteotoxicity ^26^.

**Figure 1.**
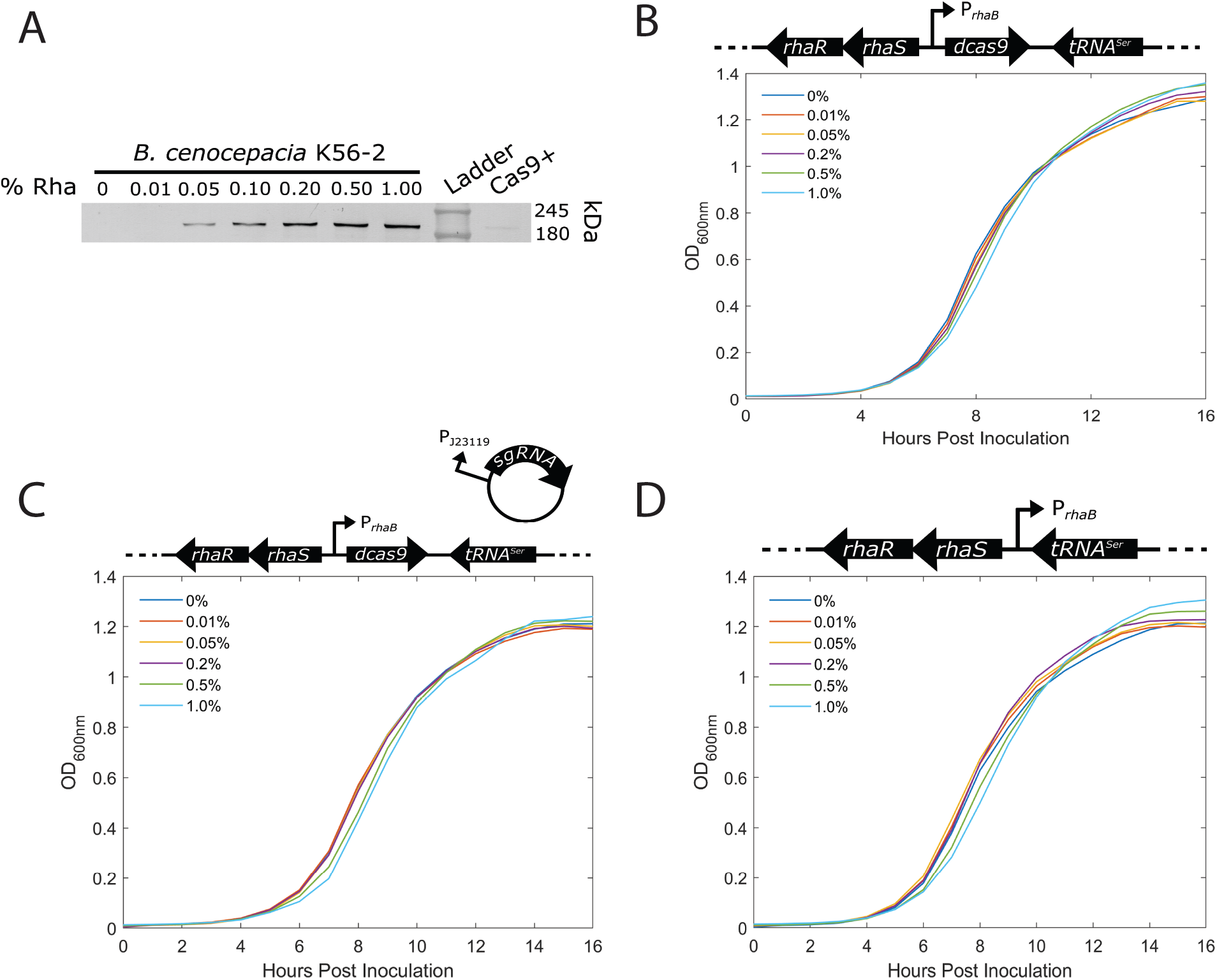
Development of CRISPRi in *B. cenocepacia* K56-2. A) dCas9 was expressed from the chromosome of K56-2 under a rhamnose inducible promoter. Cells were grown to OD_600nm_ of 0.6 then induced with rhamnose for three hours. The soluble protein fraction was run on an 8% SDS gel. dCas9 was detected by an α-dCas9 antibody followed by a second antibody linked to alkaline phosphatase. The lane labelled Cas9+ was loaded with 5 ng of purified Cas9. B, C and D) Growth curves of *B. cenocepacia* K56-2∷dCas9 (B), dCas9 expressed with a non-genome targeting sgRNA (dCas9/pgRNA-non target) (C) and K56-2∷CTX1-rha, the vector control plasmid for the integration (D) in LB media show that expression of the chromosomally-encoded dCas9 induced with rhamnose up to 1% does not affect growth.

**Figure 2.**
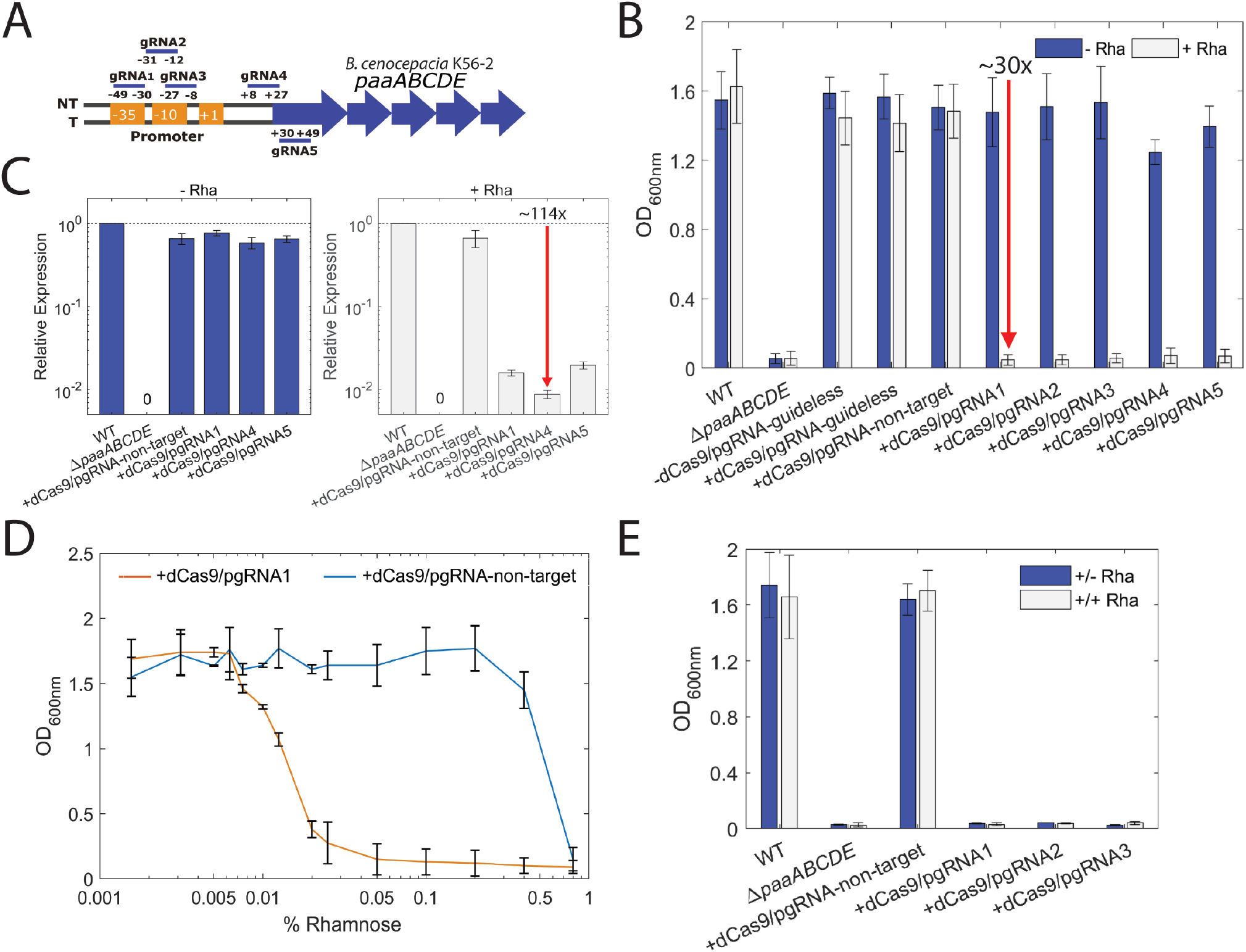
Targeting *paaA* with CRISPRi effectively suppresses growth in phenylacetic acid as the sole carbon source in *B. cenocepacia* K56-2. A) Positions of the sgRNAs targeting different regions upstream of and on *paaA*. sgRNAs 1, 2, and 3 were designed to target the promoter elements (−35 and −10 boxes) on the non-template (NT) strand. gRNA4 targeted the start codon on the NT strand and gRNA5 targeted the downstream region adjacent to the start codon on the T strand. B) CRISPRi blocks transcription in a strand non-specific manner. WT, a mutant of the *paaABCDE* operon (Δ*paaABCDE*), K56-2∷CTX1-rha (−dCas9) and K56-2∷dCas9 (+dCas9) harboring pgRNA with or without specific gRNAs were grown for 24 hours in minimal medium with PA (M9+PA) without (-Rha) or with 0.2% rhamnose (+Rha). C) RT-qPCR revealed a ~114-fold reduction in paaA mRNA, demonstrating a robust knockdown of *paaA* expression in K56-2. D) Expression of the dCas9 can be controlled by varying the amount of inducer added to the medium, providing tunability to the CRISPRi system. However, high level induction of dCas9 with rhamnose (0.4% and beyond) was lethal for the non-genome targeting mutant (dCas9/pgRNA-non-target) expressing dCas9. E) Cells were grown overnight in LB medium with 0.2% rhamnose to induce expression of dCas9. Then cells were transferred to M9+PA and grown for 24 hours with (+/+ Rha) and without (+/− Rha) 0.2% rhamnose. All the values are the average of three independent biological replicates; error bars represent arithmetic mean ± SD.

A single chromosomal copy of *dcas9* may provide sufficient levels of expression as observed previously ^18,22,27^. Using the mini-CTX system, we introduced a single copy of *dcas9* under control of the rhamnose-inducible promoter into *B. cenocepacia* K56-2 (Supplemental Figure 3A and B). We observed titratable dCas9 expression at various levels of rhamnose by immunoblot (Figure 1A and Supplemental Figure 4). At rhamnose concentrations up to 1% there was no growth defect in K56-2∷dCas9 (Figure 1B) or K56-2∷dCas9 with non-genome targeting sgRNA (pgRNA-non-target) (Figure 1C) compared to the vector control mutant (Figure 1D).

**Figure 3.**
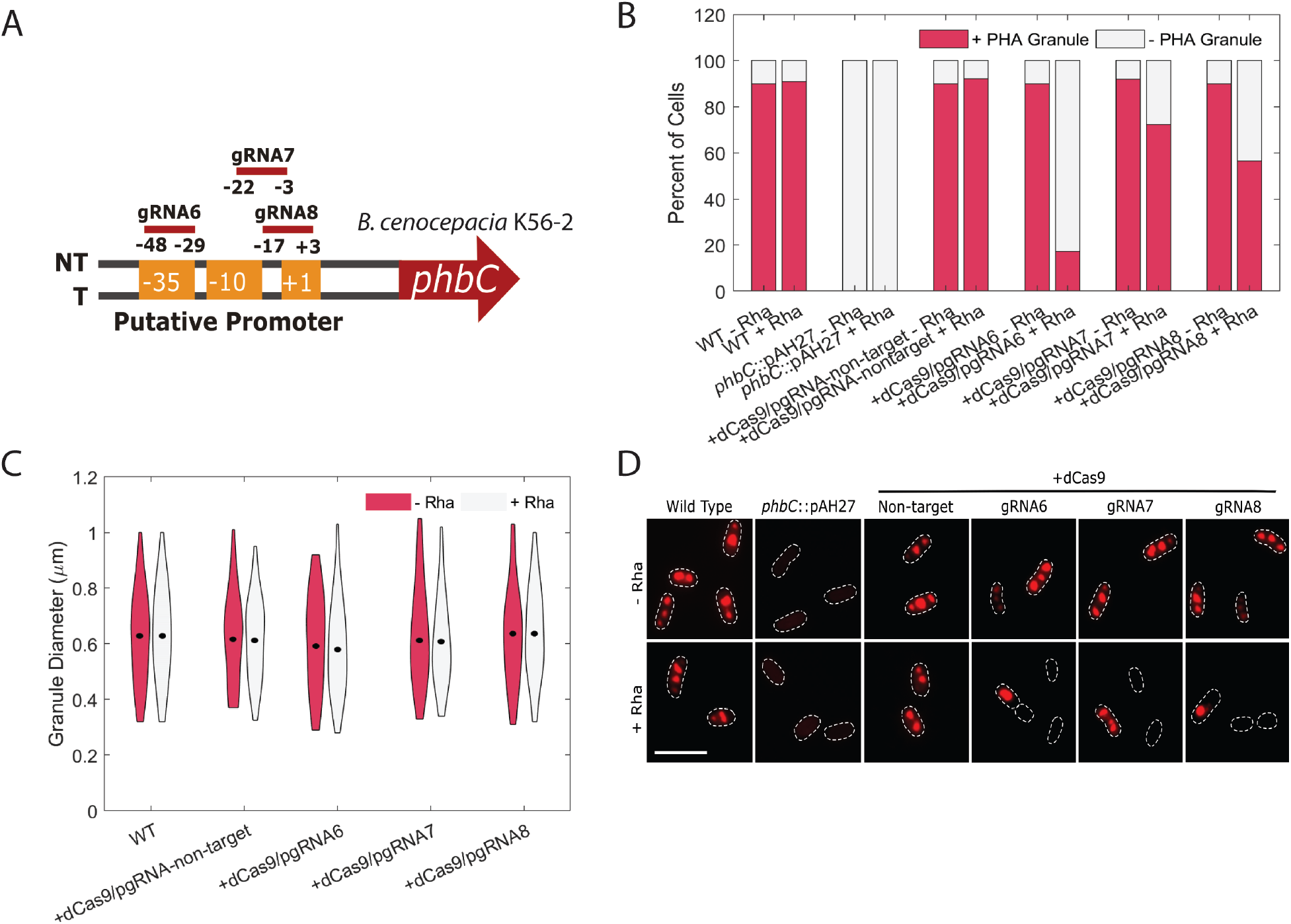
Targeting *phbC* with CRISPRi reduces polyhydroxyalkanoate (PHA) granule accumulation in K56-2. A) Positions of the sgRNAs targeting different regions upstream of *phbC*. sgRNAs 6, 7, and 8 were designed to target the promoter elements (−35 and −10 boxes) on the non-template (NT) strand. B) CRISPRi reduces but does not completely abrogate the number of cells with PHA granules. WT, an insertional mutant of the *phbC* gene (*phbC∷*pAH27), K56-2∷CTX1-rha (−dCas9) and K56-2∷dCas9 (+dCas9) harboring pgRNA with or without specific gRNAs were grown overnight without (−Rha) or with 0.2% rhamnose (+Rha). Cells were washed and stained with Nile Red, and observed by fluorescence microscopy. One to two-hundred cells were counted and the % of cells with PHA granules was calculated. C) PHA granules that remain are identical to those in the WT. Strains were grown and processed as for B and the diameter of the PHA granules was measured. Thicker areas of the violin bars represent more granules with that diameter.The mean in each condition is shown by a black dot. D) The strains were grown and processed as for B). Dashes indicate cell boundaries and the scale bar is 5 μm.

**Figure 4.**
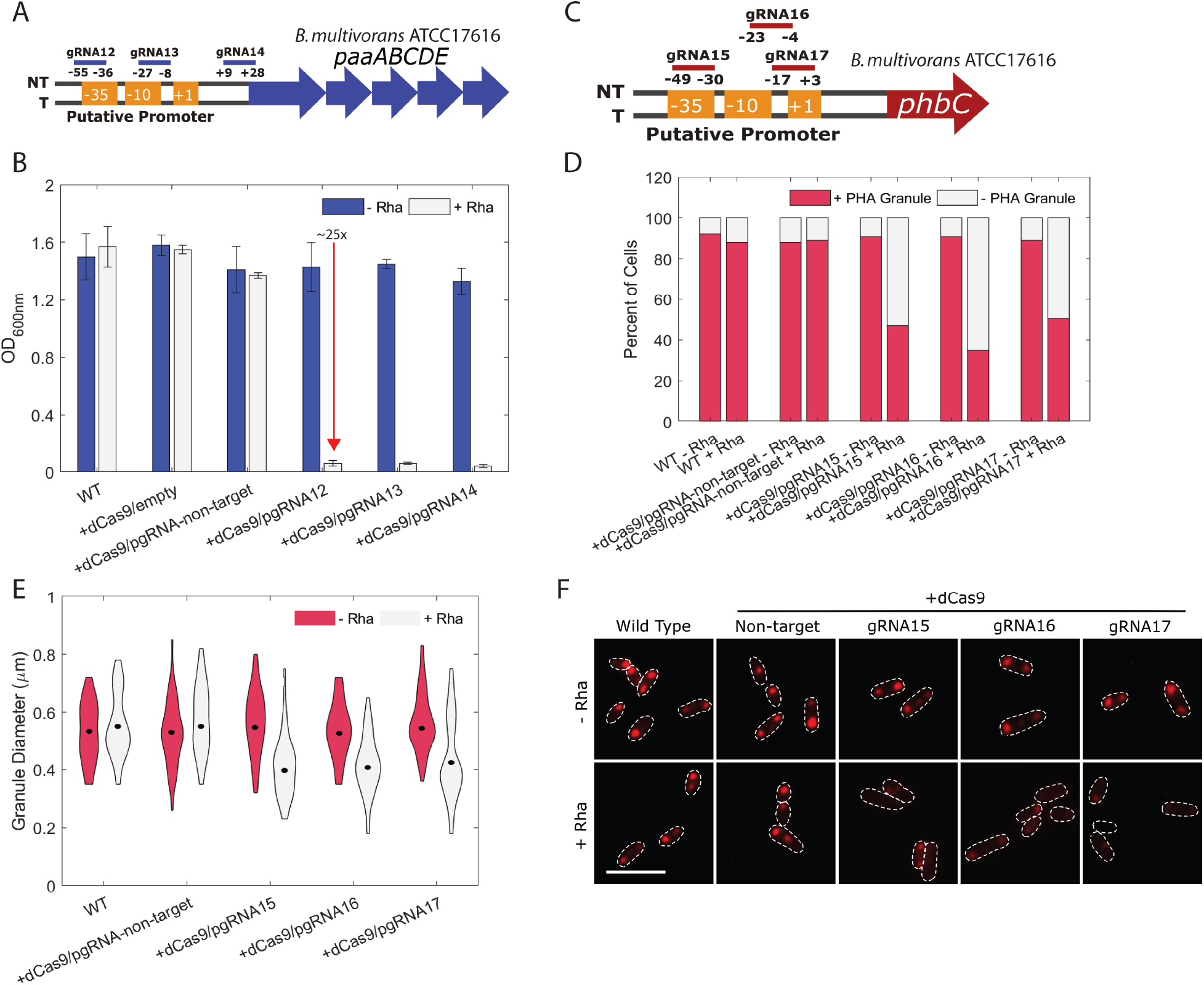
CRISPRi in *B. multivorans* ATCC17616 effectively represses *paaA* and *phbC*. A) Positions of the sgRNAs targeting upstream regions of *paaA*. sgRNAs12 and 13 were designed to target the −35 and −10 boxes of the promoter, while sgRNA14 targeted the 5’ region of the ORF. All sgRNAs targeted the NT strand. B) Targeting *paaA* suppressed growth on PA as a sole carbon source. WT, ATCC 17616∷dCas9 (+dCas9) with or without sgRNAs in A) were grown for 24 hours in M9+PA without (−Rha) or with 0.2% rhamnose (+Rha). C) Positions of the sgRNAs targeting upstream regions of *phbC*. All sgRNAs were designed to target the −35 or −10 elements of the promoter on the NT strand. D) Targeting *phbC* reduces the overall number of cells with PHA granules. Strains were grown overnight without (−Rha) or with 0.2% rhamnose (+Rha). Cells were washed and stained with Nile Red, and observed by fluorescence microscopy. One to two-hundred cells were counted and the % of cells with PHA granules was calculated. E) PHA granules that remain are smaller than those in the WT. Strains were grown and processed as for D); however, the diameter of the PHA granules was measured, with thicker areas representing more granules with that diameter. The mean in each condition is shown by a black dot. F) The strains were grown and processed as for D). Dashes indicate cell boundaries and the scale bar is 5 μm.

### Tunable and durable CRISPRi silencing of paaA suppresses growth on phenylacetic acid

To evaluate the utility of the CRISPRi system for gene repression in *B. cenocepacia* K56-2, we first chose to target the *paaA* gene, which encodes phenylacetate-CoA oxygenase subunit PaaA ^28^. This gene, and the rest of the *paaABCDE* operon, enable growth with phenylacetic acid (PA) as a sole carbon source in *B. cenocepacia* K56-2 ^29^, with the lack of growth being a clearly observable phenotype when the *paaA* gene is disrupted. In addition, as this is the first characterization of CRISPRi in *Burkholderia*, we also wished to assess the effect on repression efficiency when targeting the non-template (NT) and template (T) strands, as previous studies have demonstrated profound differences ^9,30^. We therefore designed five sgRNAs: three sgRNAs targeted the promoter elements and adjacent to the transcription start site (TSS) on the NT strand (sgRNA 1, 2 and 3), one sgRNA targeted near the start codon of *paaA* on the NT strand (sgRNA 4), and one sgRNA targeted near the start codon of *paaA* on the T strand (sgRNA 5) (Figure 2A).

For phenotypic characterization of the *paaA* mutants harbouring the codon-optimized *dcas9*, we used M9 minimal medium with PA (M9+PA) as the sole carbon source. Upon induction of dCas9, the growth of all mutants (except the controls) was suppressed approximately 30-fold (Figure 2B) to the same level of a Δ*paaABCDE* mutant, which is unable to utilize PA as a sole carbon source ^29^. We observed only moderate repression (up to ~6-fold) when using the native *dcas9* (Supplemental Figure 5). Furthermore, in the absence of rhamnose all of the mutants grew at or near wild-type levels, suggesting that *dcas9* expression is tightly repressed in non-inducing conditions. Phenotypically, we did not observe a differential effect of placement of the sgRNA-binding site, as the growth of all mutants was suppressed equally. We also found there were no differences in control strains; mutants expressing dCas9 and either a guideless or non-targeting sgRNA displayed the same levels of growth. However, RT-qPCR demonstrated that while growth was suppressed equally in the mutants, there were sgRNA-dependent differences in gene expression levels (Figure 2C). At 0.2% rhamnose, sgRNA4, targeting near the start codon of *paaA* on the NT strand, was the most effective in repressing *paaA* mRNA expression (at least 114-fold repression), whereas sgRNA1 and sgRNA5 only had ~63-fold and ~51-fold repression, respectively (Figure 2C). In the absence of rhamnose there was no difference in gene expression levels between the mutants and wild type confirming that the *dcas9* is tightly regulated in non-inducing conditions.

**Figure 5.**
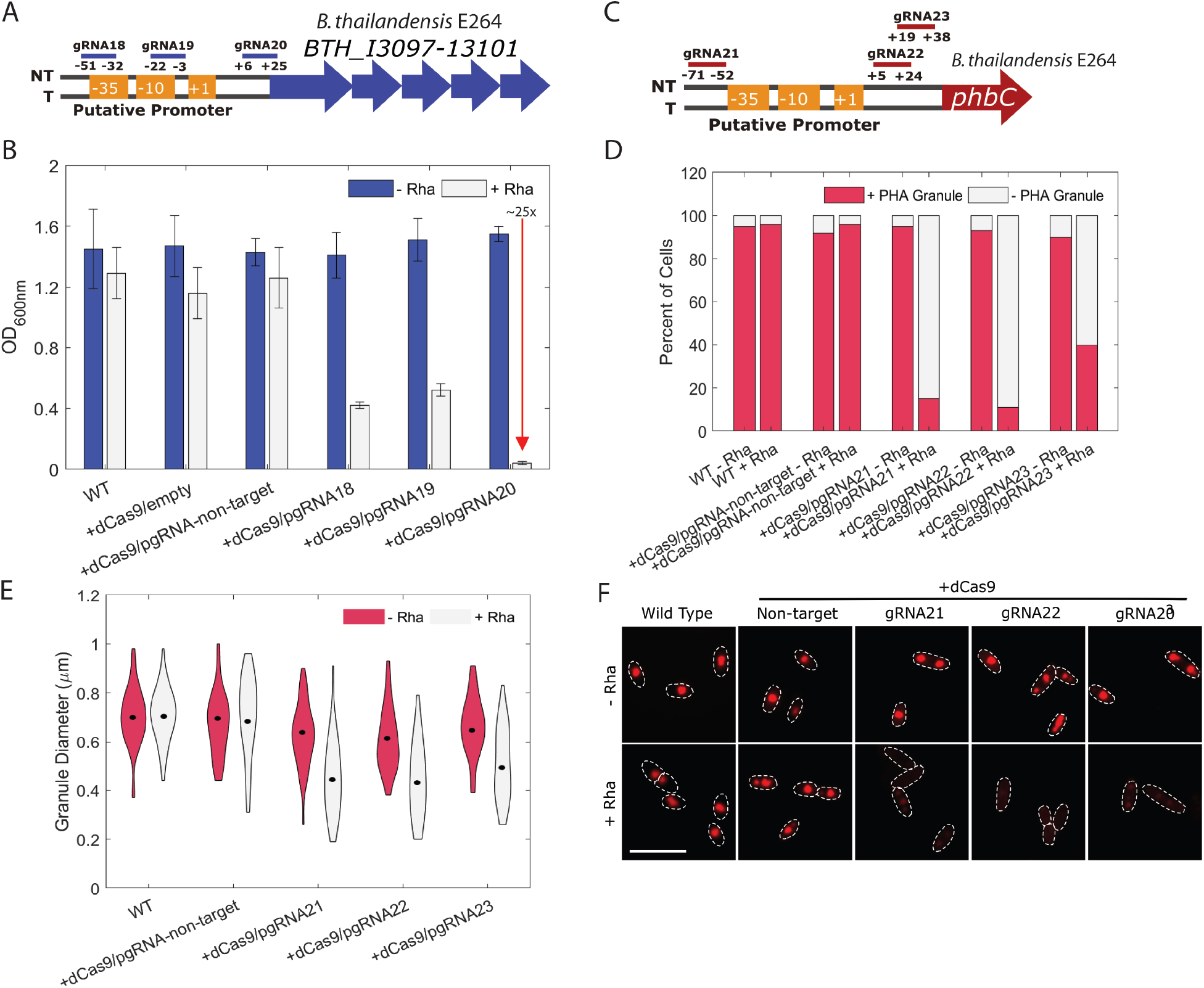
CRISPRi in *B. thailandensis* E264 effectively represses *paaA* (BTH_I3097) and *phbC*. Positions of the sgRNAs targeting upstream regions of *paaA* (BTH_I3097). sgRNAs 18 and 19 were designed to target the −35 and −10 boxes of the promoter, while sgRNA 20 targeted the 5’ region of the ORF. All sgRNAs targeted the NT strand. B) Targeting *paaA* suppressed growth on PA as a sole carbon source. WT, E264∷dCas9 (+dCas9) with or without sgRNAs in A) were grown for 48 hours in M9+PA without (−Rha) or with 0.2% rhamnose (+Rha). C) Positions of the sgRNAs targeting regions of *phbC*. sgRNA 21 targeted just upstream of the −35 box, sgRNA22 and 23 targeted just downstream of the −10 box, all on the NT strand. D) Targeting *phbC* reduces the overall number of cells with PHA granules. Strains were grown overnight without (-Rha) or with 0.2% rhamnose (+Rha). Cells were washed and stained with Nile red, and observed by fluorescence microscopy. One to two-hundred cells were counted and the % of cells with PHA granules was calculated. E) PHA granules that remain are smaller than those in the WT. Strains were grown and processed as for D); however, the diameter of the PHA granules was measured, with thicker areas representing more granules with that diameter. The mean in each condition is shown by a black dot. F) The strains were grown and processed as for D). Dashes indicate cell boundaries and the scale bar is 5 μm.

Next, we sought to determine the tunability of our CRISPRi system in *Burkholderia*. Tuning is useful to control the level of transcriptional inhibition when precision is required, such as in the study of essential genes. To that end, we examined the growth of the dCas9 mutant of *B. cenocepacia* K56-2 with the sgRNA1 (targeting *paaA*), using PA as a sole carbon source in the presence of various concentrations of rhamnose. The results showed that our CRISPRi system is tunable, exhibiting growth reduction in a dose-dependent manner with variable repression across sub-saturating rhamnose concentrations (between 0.005% and 0.05% rhamnose) (Figure 2D). This trend was confirmed by RT-qPCR (Supplemental Figure 6). We observed a nearly 30-fold repression in OD_600nm_ at concentrations well below maximum induction as identified by immunoblot, suggesting our system produces more dCas9 than is required for maximum repression as observed by this phenotype. Contrary to what had been observed in rich medium (Figure 1B and C), all dCas9 mutants (with or without the sgRNA) showed a growth defect in M9+PA above 0.2% rhamnose (Figure 2D). A similar phenomenon was also seen in M9+glycerol (data not shown). Intermediate levels of growth were observed in concentrations of rhamnose between 0.005% and 0.05%. For consistency, we therefore used 0.2% rhamnose for dCas9 induction in all experiments.

Although once rhamnose is removed from the culture, expression of dCas9 is no longer induced, it remained possible that the dCas9 already synthesized may persist and cause long-term, or durable, silencing. To address this, the mutant strains harbouring sgRNAs targeting *paaA*, were grown overnight in rich medium with 0.2% rhamnose, effectively priming the cells with dCas9. Strikingly, when grown in M9+PA with and without rhamnose, we again observed strong repression of growth (~30-fold) in all conditions regardless of the presence of rhamnose in the M9+PA, and at levels similar to the Δ*paaABCDE* mutant (Figure 2E). This suggests that after the inducer is removed, dCas9 is either slowly degraded in K56-2 or is present at high enough levels in the cells for durable repression in growth inhibiting conditions. Moreover, the lack of growth is not simply due to a loss in cell viability, as CFU/mL of the cultures did not decrease over time (Supplemental Figure 7). We likely observed a durable knockdown in this scenario due to the conditional essentiality of *paaA*. Due to the strong interaction of the dCas9:sgRNA complex with its target, repression cannot be easily relieved except by the DNA polymerase machinery (Jones et al. 2017). When transferred to medium with PA as the sole carbon source, cell growth would halt from lack of a carbon source, therefore ensuring *paaA* expression remained repressed.

### Single-cell analysis reveals an ‘all or none’ effect in B. cenocepacia K56-2

While at the culture level the effect of CRISPRi is tunable, we further explored the effect of maximum dCas9 induction at the single-cell level. We therefore targeted *fliF*, a gene encoding a transmembrane protein that forms both the S and M rings (MS ring) of the basal body complex of the flagellum ^31^. Silencing *fliF* should result in non-flagellated cells as FliF is required for flagellum formation ^32^. We targeted *fliF* by introducing four sgRNAs designed to bind near the putative promoter and start codon on the NT strand (Supplemental Figure 8A). Our goal was to assess individual cell flagellation and compare it with swimming motility at the population level. While we observed an approximately 5-fold reduction in swimming motility compared to controls in a plate-based assay (Supplemental Figure 8B and C), we were unable to observe flagella in any of the mutants harbouring gRNAs targeting *fliF* (Supplemental Figure 8D). It is possible that interfering with *fliF* expression rendered fragile flagella that could not remained attached to the cell during the staining process, while still being functional when grown in culture ^33^. In contrast to the swimming motility of the CRISPRi mutants we confirmed that insertional inactivation of *fliF* (*fliF*∷pAH26) completely ablates swimming motility, as seen previously ^34^.

To further elucidate the effect of CRISPRi at the single cell level, we targeted *phbC*, a gene encoding poly-β-hydroxybutyrate polymerase, an enzyme required for PHB synthesis. To repress *phbC* expression, we designed three sgRNAs to target the region up to 50 bp before the start codon on the NT strand, at the putative promoter site (Figure 3A). Polyhydroxyalkanoate (PHA) granule accumulation was assessed by fluorescence microscopy with Nile Red staining after overnight induction with rhamnose. For comparison to a null phenotype, we created a *phbC* insertional mutant (*phbC*∷pAH27) which was unable to produce PHA granules (Figure 3B and D). Compared to the wild-type and non-target controls, markedly few cells harbouring sgRNAs targeting *phbC* contained PHA granules, ranging from 17-70% depending on the sgRNA (Figure 3B). sgRNA6 rendered the strongest level of repression, with only 17.1% of cells containing PHA granules in contrast to 86.9% for the wild-type (Figure 3B). Interestingly, although few cells possessed PHA granules, the granules were of identical size to those in wild-type cells, averaging 0.65μm in diameter (Figure 3C and 3D). As shown by the insertional mutant, inactivation of *phbC* ablates PHA granule accumulation; therefore, the presence of granules of the same size in the dCas9 mutants as in the wild-type suggests an ‘all-or-none’ effect in *B. cenocepacia* K56-2, where most cells display the silenced phenotype, but some manage to at least temporarily escape the effect of CRISPRi.

### Broad host range of the mini-CTX system extends applicability to other species

Having shown that the *S. pyogenes* dCas9 renders strong gene repression in *B. cenocepacia* K56-2, we next turned our attention to other important species of *Burkholderia*. We introduced the codon-optimized *dcas9* gene into the chromosome of *B. multivorans* ATCC 17616 and targeted the *paaA* and *phbC* genes by designing three sgRNAs for each centered around the putative promoter (Figures 4A and C). Targeted repression of *paaA* by each of the three sgRNAs strongly suppressed growth in M9+PA by approximately 25-fold (Figure 4B), very similar to the activity observed in *B. cenocepacia* K56-2. However, we were unable to detect robust expression of dCas9 by immunoblot in *B. multivorans* ATCC17616 (Supplemental Figure 9A) despite the observed phenotype being indicative of expression of dCas9 and effective gene silencing. Expression of dCas9 in the presence of the non-targeting sgRNA did not affect growth in *B. multivorans* ATCC 17616 (Supplemental Figure 9B). Following this, targeting *phbC* rendered relatively poor inhibition of PHA granule accumulation with between 35-50% of cells containing PHA granules, depending on the sgRNA (Figure 4C and 4D). In contrast to what was seen in *B. cenocepacia* K56-2, we observed granules of reduced size (Figure 4E and 4F).

Members of the *B. pseudomallei* group are phylogenetically distinct from the Bcc ^1^, but remain of interest due to the ability to cause infection (such as melioidosis and glanders) and for their remarkable capacity for secondary metabolite production ^35^. *B. thailandensis* is a commonly used model for the pathogenic members of the *B. pseudomallei* group, we therefore also introduced the codon-optimized *dcas9* gene into the chromosome of *B. thailandensis* strain E264. When induced with rhamnose, dCas9 was highly expressed (Supplemental Figure 9A) and did not impair growth in the presence of a non-targeting sgRNA (Supplemental Figure 9B). Similarly, as for *B. cenocepacia* K56-2 and *B. multivorans* ATCC 17616, we designed three sgRNAs targeting the putative promoters of the *paaA* and *phbC* genes (Figure 5A and C). Upon induction of dCas9 with 0.2% rhamnose, growth of the mutants harbouring the *paaA*-targeting sgRNAs was suppressed in M9+PA to varying levels ranging from 3-fold (sgRNA19) to 25-fold (sgRNA20) (Figure 5B). Lastly, for the sgRNAs targeting *phbC*, the results mirror those seen in ATCC 17616, as depending on the sgRNA there was variation in the percent of cells with PHA granules (10-40%) and the diameter of the granules (overall decrease in size) (Figure 5C-F). Though the exact genetic contributions to PHA synthesis in E264 remain unclear, it has been previously suggested that PhbC is not the only polyhydroxyalkanoate polymerase in E264 ^36^.

The integration vector we implemented to deliver *dcas9* to the chromosome of various species of *Burkholderia* relies on the expression of the φCTX integrase and recombination using the plasmid-borne *attP* site with the chromosomal *attB* site ^24^. φCTX is a *Pseudomonas*-infecting phage, and the mini-CTX integration system was originally designed for use in *P. aeruginosa* ^24^. The mini-CTX integration system has been used successfully in other species; however, this has been mostly limited to members of *Burkholderia* ^37,38^. The utility in *Burkholderia* has been comparable to *P. aeruginosa*, in part owing to efficient integration. In the species used in this study, we observed the integration efficiencies to be 6×10^−7^ in K56-2, 6×10^−8^ in E264, and 5×10^−9^ in ATCC 17616 (Supplemental Figure 10A), compared to 10^−7^ to 10^−8^ observed previously in *P. aeruginosa* ^24^. To further broaden the scope of the applicability of our CRISPRi system, we used NCBI BLAST to search all published genomes for putative *attB* sites. While the full-length *attB* site is 30 nt, integration is known to occur if only the 5’ 19 nt are completely complementary, such as for many species of *Burkholderia*. This shorter *attB* site was therefore used as a BLAST query, resulting in 1760 hits with 100% alignment (Supplementary Table 5). Enterobacteria were excluded from the search as the pMB1 *oriR* in the mini-CTX system is functional is these species; therefore, integrants cannot be easily isolated. Furthermore, the search parameters were modified to only include species of Proteobacteria, as there were few hits of non-Proteobacterial species with 100% alignment (data not shown). Overall, there were 168 unique species from 67 genera. As expected, the most abundant hit corresponded to species and strains of *Pseudomonas* (480 hits), then followed by *Acinetobacter* (443 hits), *Burkholderia* (276 hits), *Neisseria* (170 hits), and *Ralstonia* (146 hits), with members of the other 62 genera comprising the remaining 245 hits. A summary of the major hits and species of interest (pathogenic, environmental, biotechnological, etc.) can be found along with the genomic context in Table 1. We note that the hit table comprises species with both high and low GC-content genomes, and while the GC-rich codon-optimized *dcas9* (in pAH-CTX1-rhadCas9) may be better suited for species with high GC-content genomes, such as those in the families *Pseudomonadaceae* and *Alcaligenaceae*, the native *dcas9* (in pAH-CTX1-rhadCas9-native) may have better functionality in species with low GC-content genomes, such as those in the families *Moraxellaceae* and *Neisseriaceae*.

**Table 1.**
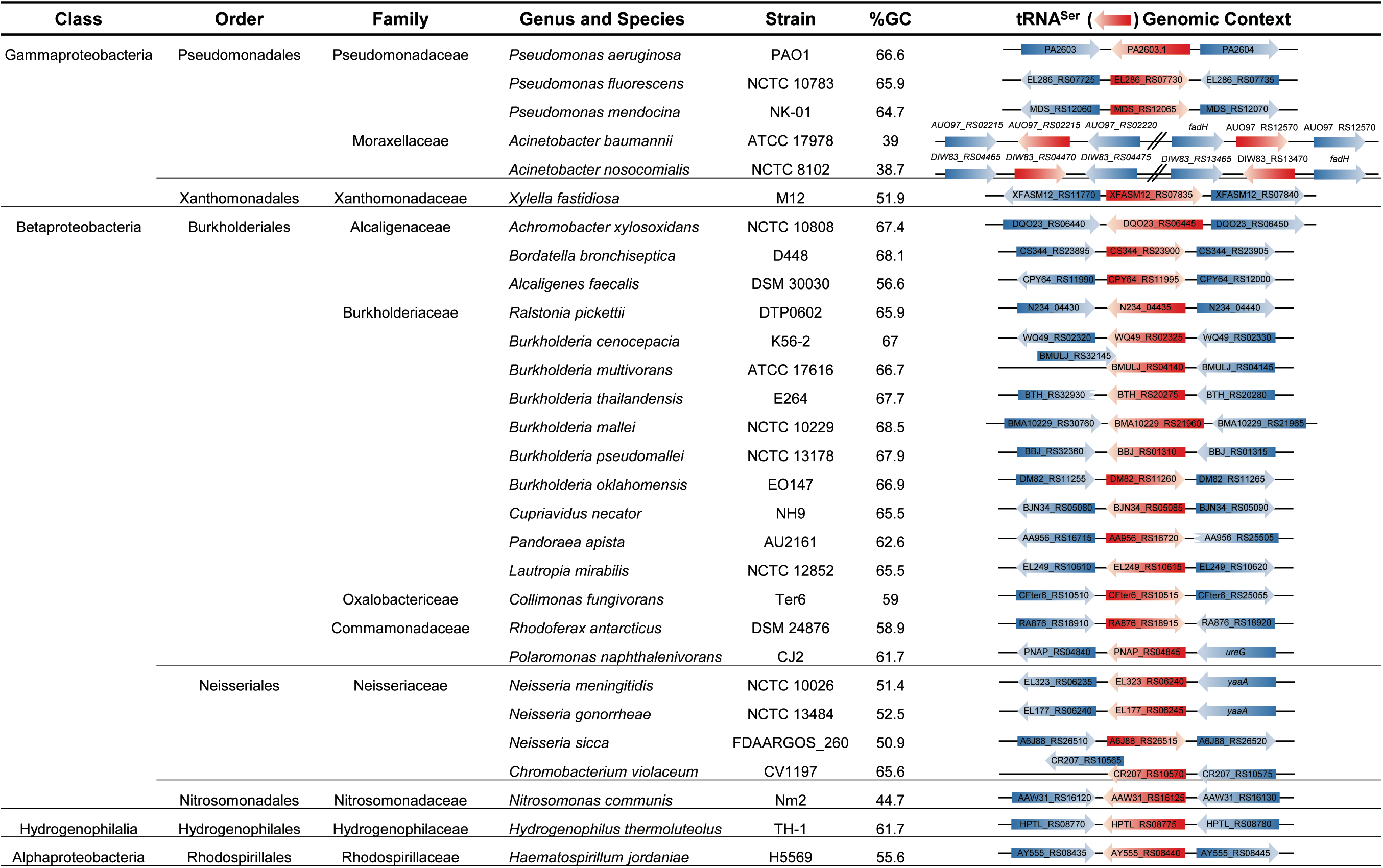
Putative host range of the mini-CTX system and genomic context of selected hit sites

A further consideration for use of our CRISPRi system is that the pFLPe recombinase-expressing plasmids might not be functional in all species. In which case, the *int* gene along with the *oriT*, pBM1 *oriR*, and the *tet* gene would not be excised. Continued expression of the integrase could cause instability of the integrated DNA and loss of the dCas9 gene over time. To explore this possibility, we assessed the stability of the mini-CTX integration in K56-2, ATCC 17616, and E264 in serial passages over four days without tetracycline selection. For all species over the entire experiment we determined CFU counts on agar with and without tetracycline selection and found equal recovery of tetracycline-resistant colonies as total colonies (Supplemental Figure 10B-D), suggesting the integration containing the dCas9 is stable.

## Discussion

Genetic tools are necessary to dissect the molecular mechanisms governing cellular processes. Here, we report the development of a CRISPRi system for efficient repression of gene expression in *Burkholderia*. By mobilizing the *dcas9* gene that was codon-optimized for the GC-rich *Burkholderia* on a broad host range mini-CTX1 integration vector, we demonstrate robust, tunable, and durable repression of endogenous genes. While others have shown effective repression using the native *dcas9* gene in *E. coli* ^9^, *B. subtilis* ^18^, *Staphylococcus aureus*, and *Acinetobacter baumannii* ^22^, codon-optimization was a necessary step for K56-2, as expression of the native *dcas9* gene was not detectable in K56-2. Indeed, for expression in species with high GC-content, codon optimization appears to be necessary. In *P. aeruginosa* (66.3% GC) both the *S. pyogenes* dCas9 ^22^ and the *S. pasteurianus* dCas9 ^13^ were not expressed unless first codon-optimized, albeit for *Homo sapiens* (optimized at 50.2% GC) or *Mycobacterium* (~67% GC), respectively. Additionally, the *S. thermophilus* dCas9 was codon-optimized for *Mycobacterium* for efficient expression in *M. tuberculosis* and *M. smegmatis* ^26^.

Upon codon-optimizing the *dcas9* gene, we observed a severe growth defect when expressed from a multicopy pBBR1-origin plasmid. At this time, we are unsure if this was caused by a metabolic burden of expressing a large protein from a multicopy plasmid or from direct proteotoxicity. Previous studies have demonstrated that expression of the canonical *S. pyogenes* dCas9 causes toxicity in *M. smegmatis*, *M. tuberculosis* ^26^, and *E. coli* ^39^, which provides rationale for developing a system with low-enough levels of dCas9 expression to maintain cell viability without sacrificing repression activity. While this effort has spurred the exploration of alternative dCas9 orthologues ^26^, we found that introducing *dcas9* in single copy in the chromosome provided a balance of repression activity without affecting growth. Furthermore, while other systems display up to 3-fold repression in the absence of inducer ^13,18^, our application of the tightly regulated rhamnose-inducible promoter from *E. coli* did not display a leaky phenotype in any of the three species tested. While we did observe increased dCas9 expression at rhamnose concentrations of 0.5% and 1.0% with no inhibitory effect on growth in LB (Figure 1C and D), the increased level of dCas9 expression was accompanied by a substantial growth defect in M9 minimal media with PA (Figure 2D) or glycerol (data not shown) as the sole carbon source. As the growth rate is decreased in minimal media, it is possible that growth inhibitory, off-target effects that are alleviated in fast growing cells by dilution of dCas9 bound DNA sites with newly replicated ones, are more evident in slowly growing cells. While the RNA polymerase is largely incapable of displacing the dCas9:sgRNA complex, the DNA replication machinery is not blocked by dCas9. In fact, it has been observed that the dissociation of the dCas9:sgRNA complex correlates well with the generation time in *E. coli* ^40^. This is an important consideration for understanding the durability of growth suppression we observed when using PA as a sole carbon source. Cells primed by pre-incubation with rhamnose are unable to grow in M9+PA, creating a condition where *paaA* is an essential gene and its absence would arrest cell division and replication. Hence, the DNA replication machinery would not displace the dCas9:sgRNA from the *paaA* gene, resulting in long-term growth suppression. By extension, we predict that strong knockdown of any (conditionally) essential gene that causes a halt in DNA replication would be durable in this manner. This is a useful aspect of CRISPRi that we are investigating further.

Although it has been reported that the CRISPRi system is less effective in silencing gene expression when the template strand (T) is targeted compared to the non-template (NT) strand, our results demonstrated nearly 30-fold suppression of growth irrespective of the *paaA* target strand (Figure 2B). This suggests that the efficiency of the CRISPRi system might not be strand specific at all loci, supporting the findings from Guo et al. ^41^. However, this effect might have been masked by the strong repression we observed, as it is difficult to compare across null phenotypes. Our findings are also supported by the RT-qPCR that demonstrated differential repression of gene expression based on the targeted location. Interestingly, our data suggests that, for *paaA*, targeting near the start codon of the NT strand (gRNA4) was more effective than targeting near the start codon of the T strand (gRNA5). Additionally, sgRNAs targeting different regions of the 5’ region of *paaA* produced a strong 25 to 30-fold repression in the three species studied here. Therefore, the observed knockdown efficiency appears to be largely unaffected by sgRNA placement on targeting regions overlapping or adjacent to the −35 and −10 promoter elements, at least for phenotypes where complete repression of gene expression is not required. The cause of this is likely due to the large dCas9:sgRNA complex that when bound to the promoter sterically interferes with RNA polymerase binding. Indeed, the ~160 kDa dCas9 enzyme has an average DNA footprint of 78.1 bp, much larger than most promoter regions ^42^. This could explain why Cui et al. ^19^, and our results for *paaA*, demonstrate strong repression when the template strand is targeted a short distance downstream of the transcription start site.

Single cell level analysis of our system suggested a unimodal ‘all or none’ effect in K56-2 with ~17% cells escaping the silenced phenotype when targeting *phbC* compared to an insertion mutant, which did not possess any PHA granules. The PHA granules in these cells were of identical diameter to wild type. The ‘all or none’ effect might be attributed to an uneven distribution of sugar transporters after cell division as observed previously for arabinose-inducible expression ^43^. If a similar mechanism involved transporting rhamnose into the cell, some cells could escape the rhamnose-mediated *dcas9* induction to silence the target gene. We hypothesize any escape would be temporary as the transporter would be expressed in growing cells, allowing rhamnose uptake. However, a rhamnose transporter in *Burkholderia* has so far not been described. Interestingly, targeting *phbC* in *B. multivorans* ATCC17616 and *B. thailandensis* E264 resulted in a mixed phenotype with a reduced number of cells with granules as well as an overall decrease in granule size. Even though some of the cells appeared to have escaped the repression (up to 35% in ATCC17616 and 6% in E264), more than 90% of the cells contained granules of reduced size. This suggests that there is an unelucidated secondary PHA synthesis pathway in *B. thailandensis* E264 and *B. multivorans* ATCC17616. Funston et al. ^36^ have observed similar results in *B. thailandensis* E264 where transposon mutants in *phb* genes (*phbA*, *phbB* and *phbC*) retained the ability to synthesize PHA, albeit at lower levels.

One of the hallmarks of CRISPRi is the broad-range amenability in diverse bacteria, enabling synthetic biology and mechanistic investigations into many dozens of species in innovative ways. We wished to apply our CRISPRi system in this manner and therefore mobilized both the native *dcas9* gene, suitable for low/medium GC-content organisms, and the *dcas9* gene, codon-optimized for GC-rich *Burkholderia*, on the mini-CTX1 integration vector. Our analysis of putative hosts (with *attB* sites near the 3’ end of the serine tRNA) identified 168 unique species in 67 genera, mostly from the β- and γ-Proteobacteria. Previous works have mobilized *dcas9* on broad host-range integrative plasmids ^13,22^; however, both systems use the mini-Tn*7* system, which has multiple insertion locations in certain genomes, such as many species of *Burkholderia*. Together, our work contributes to the available genetic toolkit for rapid functional analysis of bacteria.

## Methods

### Strains, selective antibiotics, and growth conditions

All strains and plasmids are found in Supplemental Table 1. All strains were grown in LB-Lennox medium (Difco). *B. cenocepacia* K56-2 and strains of *E. coli* were grown at 37°C, while *B. thailandensis* E264 and *B. multivorans* ATCC 17616 were grown at 30°C. The following selective antibiotics were used: chloramphenicol (Sigma; 100 μg/mL for *B. cenocepacia*, 20 μg/mL for *E. coli*), trimethoprim (Sigma; 100 μg/mL for strains of *Burkholderia*, 50 μg/mL for *E. coli*), tetracycline (Sigma; 50 μg/mL for all strains of *Burkholderia*, 20 μg/mL for *E. coli*), kanamycin (Fisher Scientific; 250 μg/mL for *B. thailandensis*, 150 μg/mL for *B. multivorans*, 40 μg/mL for *E. coli*), ampicillin (Sigma; 100 μg/mL for *E. coli*), gentamicin (Sigma; 50 μg/mL for all strains of *Burkholderia*).

### Construction of pSC-rhadCas9, pAH-CTX1-rhadCas9, pAH-CTX1-rhadCas9-native, and dCas9 insertional mutants

The endogenous *cas9* gene from *S. pyogenes* has low GC content (averaging 34.1%) and subsequently poor codon usage for GC-rich organisms (http://www.kazusa.or.jp/codon/). The nuclease-inactive variant (*dcas9*) was therefore codon optimized for *B. cenocepacia* by purchasing the optimized gene in two fragments from IDT (2462 bp and 1849 bp, Supplemental Table 2), each with 38 bp overlapping regions. Strong, rho-independent terminators were added following the gene. The full-length gene (with terminal *Nde*I and *Hind*III cut sites) was synthesized by overlap-extension PCR using Q5 polymerase with high GC buffer (NEB) and primers 979 and 987 (Supplemental Table 2). The first ten rounds of PCR were performed without primers to synthesize the full-length product using the overlap regions; primers were added for the following 25 cycles. The cycle parameters are as follows: 98°C for 30 sec, (98°C for 10 sec, 67.5°C for 20 sec, 72°C for 2.5 min)x10 cycles, 72°C for 5 min, 98°C for 30 sec, (98°C for 10 sec, 62.5°C for 20 sec,72 °C for 2.5 min)x25 cycles, 72°C for 10 min. The 4267 bp product was gel-purified (Qiagen) and introduced into pSCrhaB2 by restriction cloning using *Nde*I and *Hind*III (NEB). The resulting plasmid, pSC-rhadCas9, was transformed into *E. coli* DH5α, and trimethoprim-resistant colonies were screened by colony PCR with primers 954 and 955. Triparental mating with *E.* coli MM290/pRK2013 as a helper was performed as previously described ^44^.

To introduce the optimized *dcas9* into the mini-CTX1 insertion plasmid ^24^ serial restriction cloning was used (Supplemental Figure 3A). Briefly, the rhamnose-inducible promoter from pSC201 ^45^ was first PCR amplified with Q5 polymerase and primers 976 and 1071, containing *Hind*III and *Spe*I restriction sites, respectively. This fragment was introduced into mini-CTX1, to create pAH-CTX1-rha, and tetracycline-resistant *E. coli* DH5α were screened by colony PCR using primers 957 and 1074. The fragment containing the dCas9 gene was PCR amplified as above, but instead using primers 1072 and 1073, introducing *Spe*I and *Not*I restriction sites, respectively. This fragment was introduced into pAH-CTX1-rha, to create pAH-CTX1-rhadCas9, and tetracycline-resistant *E. coli* DH5α colonies were screened by colony PCR using primers 954 and 955.

The native (non codon-optimized) *dcas9* was also introduced into pAH-CTX1-rha. The native *dcas9* and transcriptional terminators were PCR amplified from pdCas9-bacteria (Addgene plasmid # 44249) with Q5 polymerase (NEB) using primers 1216 and 1217. The PCR product was cloned into pAH-CTX1-rha (creating pAH-CTX1-rhadcas9-native) using *Not*I and *Spe*I restriction sites then transformed into *E. coli* DH5α. Tetracycline-resistant colonies were screened by PCR using primers 954 and 1218.

pAH-CTX1-rha, pAH-CTX1-rhadCas9, and pAH-CTX1-rhadCas9-native were introduced into *Burkholderia* species by triparental mating using *E. coli* MM290/pRK2013 as a helper as above. Tetracycline-resistant colonies were screened by colony PCR using primer 954 (for pAH-CTX1-rha) or 1075 (for pAH-CTX1-rhadCas9) or 1219 (for pAH-CTX1-rhadcas9-native) and 1008 (for *B. cenocepacia*), 1167 (for *B. multivorans*), or 1168 (for *B. thailandensis*).

To remove the tetracycline resistance and integrase genes from the insertional mutants constructed with pAH-CTX1-rhadCas9 and pAH-CTX1-rha (Supplemental Figure 3), the Flannagan method ^46^ was used for *B. cenocepacia* K56-2, while the pFLPe system ^47^ was used for *B. multivorans* ATCC 17616 and *B. thailandensis* E264. To remove the regions flanking the *FRT* sites in *B. cenocepacia* K56-2, a fragment with 475 bp overlapping the upstream and downstream regions of the *FRT* sites was designed and synthesized (IDT) with *Kpn*I and *Eco*RI restriction sites, respectively. The fragment was ligated into pGPI-*Sce*I ^46^ via the *Kpn*I and *Eco*RI restriction sites, creating pAH18, and transformed into *E. coli* SY327. Trimethoprim-resistant colonies were screened for the insertion of the fragment with primer 153 and 154. pAH18 was introduced into the mutant backgrounds via triparental mating, as described above. Trimethoprim-resistant K56-2 were screened by PCR for both possible integration orientations using primers 154 and 1126, or 153 and 1133. To initiate the second recombination, an *Sce*I-expressing plasmid is required; however, the conventional plasmid, pDAI-*Sce*I, confers tetracycline resistance and could not be selected for in the mutant background. Therefore, the tetracycline resistance cassette was removed by digestion with *Age*I and *Xho*I. The chloramphenicol resistance gene *cat* was PCR amplified from pKD3 ^48^ using primers 1084 and 1150, then ligated into the *Age*I and *Xho*I-digested pDAI-*Sce*I backbone and transformed into *E. coli* DH5α, creating pAH25-*Sce*I. Chloramphenicol-resistant colonies were screened with primers 1091 and 1150. pAH25-*Sce*I was introduced into the mutant backgrounds by triparental mating as described above. Chloramphenicol-resistant colonies were screened for sensitivity to trimethoprim (indicating excision of pAH18) and tetracycline (indicating excision of the genes between the *FRT* sites), and then screened by PCR with primers 1126 and 1133, which bridge the excision.

The pFLPe system was used to remove the tetracycline resistance and integrase genes in the dCas9 mutants in *B. multivorans* ATCC 17616 and *B. thailandensis* E264. Triparental mating to introduce pFLPe4 into the strains was performed as for K56-2 above, except 0.2% rhamnose was added to the mating and antibiotic selection plates. Tetracycline-sensitive colonies were screened by PCR using primers 957 and 1194 (for *B. multivorans*) or 1195 (for *B. thailandensis*). pFLPe4 has a temperature-sensitive origin of replication; therefore, mutants were grown overnight in LB without antibiotics at 37°C. Single colonies were then tested for kanamycin sensitivity and then by colony PCR for pFLPe4 using primers 1128 and 1129.

### Design and construction of the sgRNA-expressing plasmids

PAM sequences closest to the 5’ end of the transcription start site (TSS) were first identified on both the non-template and template strands. We extracted 20-23 nucleotides adjacent to the PAM sequence to design the base-pairing region of the sgRNAs in the following format: 5’-CCN-N_(20-23)_-3’ for targeting the non-template strand and 5’-N_(20-23)_-NGG-3’ for the template strand. To score the specificity and identify off-target binding sites, the 5’ end of the 20-23nt variable base-pairing sequences were trimmed one base at a time and the remaining base-pairing region was searched against the appropriate organism’s reference genome. This was repeated until only 10 nt were used as a search query. Potential sgRNAs were discarded if off-target sites were discovered in this manner.

The expression vector pSCrhaB2 ^25^ was chosen as the method of sgRNA expression due to the broad host range of the pBBR1 origin of replication. The sgRNA cassette from pgRNA-bacteria ^9^ (Addgene plasmid # 44251) was introduced into pSCrhaB2 by restriction cloning with *Eco*RI and *Hind*III (NEB) to create pSCrhaB2-sgRNA. To remove *rhaS* and *rhaR*, inverse PCR was performed using Q5 polymerase (NEB) and primers 847 and 1025. The resulting fragment was ligated by blunt-end ligation using 1 μL of PCR product incubated with 0.5 μL *Dpn*I, 0.5 μL T4 polynucleotide kinase, and 0.5 μL T4 ligase (NEB) with quick ligation buffer (NEB) at 37°C for 30 minutes. The resulting plasmid, pSCB2-sgRNA, was screened using primers 781 and 848, which span the ligated junction. Individual sgRNAs were introduced into pSCB2-sgRNA using inverse PCR as previously described ^9^ (Supplemental Table 3).

### Construction of insertional mutants K56-2 fliF∷pAH26 and K56-2 phbC∷pAH27

Inactivation of *fliF* was performed with the mutagenesis system of Flannagan et al. ^49^. Briefly, a 322 bp internal fragment of *fliF* was PCR amplified from the K56-2 genome using primers 1156 and 1157 and Q5 polymerase (NEB). The fragment and pGPΩ-Tp were double digested with *Kpn*I and *Eco*RI (NEB) and ligated with T4 ligase (NEB). The resulting plasmid, pAH26, was electroporated into *E. coli* SY327, and trimethoprim-resistant colonies were screened by colony PCR for the *fliF* fragment. Triparental matings were performed as above. Trimethoprim-resistant exconjugants, were screened by motility assay (below).

Inactivation of *phbC* (WQ49_RS30385) was performed as for *fliF*. Briefly, a 328 bp internal fragment of *phbC* was PCR amplified from the K56-2 genome using primers 1196 and 1197 and Q5 polymerase (NEB). The plasmid, pAH27, created from ligating the fragment into pGPΩ-Tp using *Kpn*I and *Eco*RI (NEB) restriction sites, was electroporated into *E. coli* SY327 and trimethoprim-resistant colonies were screened by colony PCR for the *phbC* fragment. Triparental matings were performed as above, and trimethoprim-resistant exconjugants were screened by staining for polyhydroxyalkanoate granule accumulation (below).

### Assays for integration efficiency and stability of the mini-CTX1-based system

To assess integration efficiency, triparental matings were started as above. However, after the mating on LB agar, the pellicles were serially diluted and plated for CFU/mL on LB agar with 50 μg/mL gentamicin and LB agar with 50 μg/mL tetracycline and 50 μg/mL gentamicin.

To assess stability of the integration, cultures of the dCas9 mutants (containing the tetracycline resistance cassette) were serial passaged over 4 days without antibiotics. Each day, a fresh culture was started with a 1:2500 dilution of the previous day’s stationary phase culture. In addition, the cultures were serially diluted and plated for CFU/mL on LB agar without antibiotics and LB agar with 50 μg/mL tetracycline.

### Growth assay with phenylacetic acid as the sole carbon source

Overnight cultures, started from isolated colonies, of the appropriate strains were washed at 4000 xg for 4 minutes and resuspended in PBS (2.7 mM KCl, 136.9 mM NaCl, 1.5 mM KH_2_PO_4_, 8.9 mM Na_2_HPO_4_, pH 7.4) to remove growth medium. The OD_600nm_ of the cultures was normalized to 0.01 in M9 medium supplemented with 5 mM phenylacetic acid, 100 μg/mL trimethoprim, and 0.2% rhamnose as required. The culture was added to wells of a 96-well plate and incubated with continuous shaking at 37°C (*B. cenocepacia* K56-2) or 30°C for *B. multivorans* ATCC 17616 and *B. thailandensis* E264). The OD_600nm_ of the cultures was measured after 24 hours for *B. cenocepacia* K56-2 and *B. multivorans* ATCC 17616, or 48 hours for *B. thailandensis* E264.

### Fluorescent microscopy and polyhydroxyalkanoate granule detection

Overnight cultures of the appropriate strains with or without rhamnose were first washed to remove growth medium and resuspended in PBS. Cells were fixed in 3.7% formaldehyde + 1% methanol at room temperature for 10 minutes (*B. cenocepacia* K56-2) or 20 minutes (*B. multivorans* ATCC 17616 and *B. thailandensis* E264) then quenched by the addition of an equal volume of 0.5 M glycine. The cells were washed and resuspended in PBS with 0.5 μg/mL Nile Red (Carbosynth) and stained at room temperature in the dark for 20 minutes, after which the cells were washed to remove excess stain and resuspended in PBS. The cells were mounted on 1.5% agarose pads and imaged by fluorescence microscopy at 1000× total magnification on an upright AxioImager Z1 (Zeiss). Nile Red was excited at 546/12 nm and detected at 607/33 nm.

### Plate-based motility assay

Assays were performed as previously described ^50^, with some modifications. Briefly, strains were grown on LB agar with the appropriate antibiotics and single colonies were stab-inoculated into motility medium consisting of nutrient broth (Difco) with 0.3% agar. Medium was supplemented with rhamnose (Sigma) as appropriate. Plates were incubated right-side up for 22 hours at 37°C.

### Flagellum staining

Staining was performed as previously described ^50^. Briefly, an overnight culture was rested statically at room temperature for 20 minutes. Gently, a 1 in 10 dilution was prepared in water and rested statically for a further 20 minutes. A small drop of the diluted culture was placed on a clean glass slide and rested for 20 minutes. A coverglass was gently applied and one side was flooded with Ryu flagellum stain (Remel), then allowed to dry for 2 hours at room temperature. Slides were observed by light microscopy at 1000x total magnification on an upright AxioImager Z1 (Zeiss).

### SDS-PAGE and Immunoblotting

Cells from an overnight culture were subcultured into fresh medium and grown at 37°C (30°C for *B. thailandensis*) to an OD_600nm_ of 0.4, then exposed to various concentrations of rhamnose for 3 hours. Soluble protein was isolated first by sonicating the cells in TNG Buffer (100 mM Tris-HCl, 150 mM NaCl, 10% glycerol, pH 7.4) then by centrifugation at 15 000g for 20 minutes. Following boiling denaturation in SDS loading buffer (50 mM Tris-HCl, 2% SDS, 0.2% bromophenol blue, 20% glycerol, 100 mM DTT, pH 6.8), samples were run on an 8% Tris/glycine gel. To ensure equal loading, 20 μg protein was loaded per well (as determined by NanoDrop) and gels were run in duplicate (one for immunoblot, and another for Coomassie staining). Protein was transferred by iBlot to a PVDF membrane, blocked in 5% skim milk-TBST (150 mM NaCl, 10 mM Tris-HCl, 0.5% Tween-20, pH 7.5) at room temperature for 1 hour, then probed with a 1:2 000 dilution of primary α-Cas9 antibody (ThermoFisher 10C11-A12) in 5% skim milk-TBST overnight at 4°C. Following washes, the blot was probed with a 1:20 000 dilution of secondary antibody linked to alkaline phosphatase (ThermoFisher G-21060) in 5% skim milk-TBST for 1 hour at room temperature. Protein was detected by incubation with a solution of NBT/BCIP (Roche) as per the manufacturer’s protocol.

### RNA extraction and Reverse Transcription quantitative PCR (RT-qPCR) analysis

Cells from an overnight culture were subcultured at an OD_600nm_ of 0.01 into fresh medium with antibiotic and rhamnose, as necessary, and grown for 8 hours. Cells were harvested by centrifugation (3 minutes at 4600xg) and pellets were stored at −80°C until RNA extraction. RNA was purified and DNAse treated using the Ribopure bacteria kit (Ambion) with extended DNAse treatement (2 hours). RNA quality was verified by running on a 2% agarose gel. cDNA was synthesized with the iScript Reverse Transcriptase kit (Bio-Rad) and qPCR was performed using iQ SYBR Green mastermix (Bio-rad) on a CFX96 Touch Real-Time PCR Detection System (Bio-Rad). Primer efficiency was determined for each primer set and efficiencies between 95% and 105% were deemed acceptable. Data was analyzed using the comparative C_T_ method ^51,52^. Genes were normalized to a commonly used reference gene, the RNA polymerase sigma factor *sigE* (BCAM0918) ^53,54^.

### Plasmid availability

The following plasmids have been deposited to Addgene for distribution: pAH-CTX1-rha (#129390), pAH-CTX1-rhadCas9 (#129391), pAH-CTX1-rhadCas9-native (#129392), pAH25-SceI (#129389), pSCB2-sgRNA (#129463), pgRNA-guideless (#129464), and pgRNA-non-target (#129465).

## Acknowledgements

This work was financially supported by grants from the Cystic Fibrosis Foundation, Cystic Fibrosis Canada, and the Natural Sciences and Engineering Research Council of Canada (NSERC) to STC; AMH was supported by grants from the Canadian Institutes of Health Research (CIHR) and Cystic Fibrosis Canada.

The authors are grateful to Eric Déziel from the Institut National de la Recherche Scientifique – Institut Armand-Frappier for providing the miniCTX1 integration vector.

