## Supplemental information for "A broad-host-range CRISPRi toolkit for silencing gene expression in *Burkholderia*" (from another version of this preprint: version 2, 2019-06-01)

**Supplemental material:**

Supplemental Tables 1-3 are provided here

Supplemental Tables 4 and 5 are provided as separated Excel files.

Supplemental Figures 1-7 are provided here

**Supplemental Table 1:** Bacterial strains and plasmids used in this work.

| Strain or plasmid | Features | Source |
|---|---|---|
| *Burkholderia. cenocepacia* K56-2 | Clinical isolate from cystic fibrosis patient; ET12 lineage | (Darling et al., 1998) |
| K56-2::CTX1-rha | Derived from K56-2; pAH-CTX1-rha integrated at *attB* site; clean deletion of plasmid accessory genes | This study |
| K56-2::dCas9 | Derived from K56-2; pAH-CTX1-rhadCas9 integrated at *attB* site; clean deletion of plasmid accessory genes | This study |
| K56-2::dCas9-native | Derived from K56-2; pAH-CTX1-rhadCas9-native integrated at *attB* site; clean deletion of plasmid accessory genes | This study |
| K56-2 *fliF*::pAH26 | Derived from K56-2; pAH26 integrated into *fliF*; $Tp^r$ | This study |
| K56-2 *phbC::*pAH27 | Derived from K56-2; pAH27 integrated into *fliF*; $Tp^r$ | This study |
| K56-2 Δ*paaABCDE* | Derived from K56-2; clean deletion of the *paaABCDE* operon | (Pribytkova et al., 2014) |
| *Burkholderia multivorans* ATCC 17616 | Soil sample | ATCC |
| ATCC 17616::dCas9 | Derived from ATCC 17616; pAH-CTX1-rhadCas9 integrated at *attB* site; clean deletion of plasmid accessory genes | This study |
| *Burkholderia thailandensis* E264 | Rice-field soil sample in Thailand | DSMZ |

| | | |
|---|---|---|
| E264::dCas9 | Derived from ATCC E264; pAH-CTX1-rhadCas9 integrated at *attB* site; clean deletion of plasmid accessory genes | This study |
| *E. coli* DH5α | $F^-$ Φ80*lac*ZΔM15 Δ(*lac*ZYA-*arg*F) U169 *rec*A1 *end*A1 *hsd*R17 ($r_H^-$, $m_K^+$) *pho*A *sup*E44 $\lambda^-$ *thi*-1 *gyr*A96 *rel*A1 | Cardona lab collection |
| *E. coli* SY327 | $F^-$ *araD* Δ(*lac-proAB*) *argE*(Am) *recA56* Rifr *nalA* λ*pir* | (Miller and Mekalanos, 1988) |
| *E. coli* MM290 | F-, φ 80 lacZΔM15 *endA1 recA1 hsdR17* ($r_H^-m_K^+$) *supE*44 *thi*-1 Δ*gyrA96* (*ΔlacZYA-arg*F) U169 *relA*1 | Cardona lab collection |
| mini-CTX1 | $Tet^r$ , Ω-FRT-*attP*MCS, $ori_{pMB1}$, $int_{\varphi CTX}$ , and oriT | (Hoang et al., 2000) |
| pKD3 | $Cm^r$ *rgnB* $ori_{R6K}$ | (Datsenko and Wanner, 2000) |
| pAH18 | Derived from pGPI-*Sce*I; $ori_{R6K}$ $Tp^r$ $mob^+$ carries I-SceI gene; 475 bp from both upstream and downstream flanking regions between the *FRT* sites on pAH-CTX1-rhadCas9 cloned between *Kpn*I and *Eco*RI sites | This study |
| pHA25-S*ce*I | Derived from pDAI-*Sce*I; $ori_{pBBR1}$ $Cm^r$ $mob^+$ P*dhfr I-SceI*; *cat* gene from pKD3 cloned into *Age*I and *Xho*I sites | This study |
| pFLPe4 | $Ap^r$ $Km^r$ *oriT* $ori_{RO1600}$-*rep*($Ts_{Bt}$) *rhaR rhaS* $P_{rhaB}$ | (Choi et al., 2008) |
| pAH26 | pGPΩ-TP with 330 bp 5' fragment of *B. cenocepacia* K56-2 *fliF* cloned between *Kpn*I and *Eco*RI sites | This study |
| pAH27 | pGPΩ-TP with 335 bp 5' fragment of *B. cenocepacia* K56-2 *phbC* cloned between *Kpn*I and *Eco*RI sites | This study |

| | | |
|---|---|---|
| pAH-CTX1-rha | Derived from mini-CTX1; *rhaS rhaR* $P_{rhaB}$ from pSC201 cloned into *Spe*I and *Hind*III sites | This study |
| pAH-CTX1-rhadCas9 | Derived from pAH-CTX1-rha; codon-optimized *dcas9* cloned into *Spe*I and *Not*I sites | This study |
| pAH-CTX1-rhadCas9-native | Derived from pAH-CTX1-rha; native *dcas9* cloned into *Spe*I and *Not*I sites | This study |
| pSCrhaB2 | $ori_{pBBR1}$ *rhaR*, *rhaS*, $P_{rhaB}$ $Tp^R$ $mob^+$ | (Cardona and Valvano, 2005) |
| pSCrhaB2-sgRNA | Derived from pSCrhaB2; sgRNA cassette from pgRNA-bacteria cloned into *Kpn*I and *Eco*RI sites | This study |
| pSCB2-sgRNA | Template for pgRNA creation by inverse PCR; derived from pSCrhaB2-sgRNA by inverse PCR to remove *rhaS, rhaR,* and $P_{rhaB}$ | This study |
| pgRNA-guideless | Derived from pSCB2-sgRNA; 20nt binding region of sgRNA removed by inverse PCE | This study |
| pgRNA-non-target | Derived from pSCB2-sgRNA; random 20nt sequence added as sgRNA binding region by inverse PCR | This study |
| pgRNA1-23 | Derived from pSCB2-sgRNA by inverse PCR to add new 20 nt sgRNA binding region; see Supplementary Table 4 for details | |
| pSC-rhadCas9 | Derived from pSCrhaB2; codon-optimized *dcas9* was cloned into *Nde*I and *Hind*III cites | This study |
| pDAI-*Sce*I | $ori_{pBBR1}$ $Tet^r$ $mob^+$ P*dhfr I-SceI* | (Flannagan et al., 2008) |
| pGPI-*Sce*I | $ori_{R6K}$ $Tp^r$ $mob^+$ carries I-SceI gene | (Flannagan et al., 2008) |
| pGPΩ-TP | $ori_{R6K}$, $\Omega Tp^r$, $mob^+$ | (Flannagan et al., 2007) |

| | | |
|---|---|---|
| pRK2013 | $ori_{colE1}$ RK2 derivative $Kan^r$ $mob^+$ $tra^+$ | (Figurski and Helinski, 1979) |
| pSC201 | $ori_{R6K}$ *rhaR rhaS* $P_{rhaB}$ *dhfr* | (Ortega et al., 2007) |

**Supplemental Table 2.** Primers used in this work.

| Primer Number | Sequence (restriction sites in lowercase) |
|---|---|
| 153 | GTGGATGACCTTTTGAATGACCTTT |
| 154 | ACAGGAACACTTAACGGCTGACATG |
| 781 | TAAGATggatccTCAGTTGGCTTCATCGCTAC |
| 847 | TTCCTGTCAGTAACGAGAAGG |
| 848 | CCGCCAGGCAAATTCTGTTT |
| 954 | GGGTTCTATCGCCACGGAC |
| 955 | ATTTTTGCGGCGCGTGTAG |
| 957 | GTCAGCAATTTCGCCAGCAG |
| 976 | TATAATAaagcttATGTCAACTGGGTTCGTGC |
| 979 | TATATAaagcttACGCAGAAAGGCCCACCCGAAG |
| 987 | AAATTAcatatgGACAAGAAGTACTCGATCGGCC |
| 1008 | GTTTCCTCAACGACCTGCAG |
| 1025 | ATAATAggtaccCGCACGAACCCAGTTGAC |
| 1071 | ATAATAactagtTCCTGCTGAATTTCATTACGACC |
| 1072 | ATAATAactagtTATGGACAAGAAGTACTCGATCGG |
| 1073 | ATAATAgcggccgcACGCAGAAAGGCCCACCCGAAG |
| 1074 | AAGTGGATCAGCAAGGACGG |
| 1075 | AAGTCCTGAGCGCCTACAAC |
| 1077 | CCGTGCCATGTTCGACAATC |
| 1084 | ATAATActcgagCGCCTACCTGTGACGGAAGA |

| | |
|---|---|
| 1091 | ACTGCCTACCCCACAACAAC |
| 1126 | TCGTTGCGCAGATAGGTCAC |
| 1128 | GATCGATCCTACCCCTTGCG |
| 1129 | GCCTAAGGTGCTTGTTCGTC |
| 1150 | AATTAAaccggtACTCATCGCAGTACTGTTGTATTC |
| 1156 | TAAGATggtaccTTCCCTACAAGTTCGCCGATG |
| 1157 | TAAGATgaattcGGGTAGAGGTCGACGAACAC |
| 1167 | CTGCCGTTCGAGTTCATGC |
| 1168 | CGGTTACAATGCTCGACCTTG |
| 1194 | AGGGCTTCATGACGGGTTTC |
| 1195 | TCGGTATCTCTGGCTGTTCG |
| 1196 | ATAATAggtaccCGTTCCAGATGCCGTCGATG |
| 1197 | ATAATAgaattcGTGAAACGGATGCGCTCAC |
| 1216 | ATAATAactagtTATGGATAAGAAATACTCAATAGGCTT |
| 1217 | ATAATAgcggccgcACGCAGAAAGGCCCAC |
| 1218 | CTTCCGCTGTCTCTCCACTG |
| 1219 | GTAAACGGATGCTGGCTAGTG |

**Supplemental Table 3.** Inverse PCR primers (to introduce sgRNAs) and colony PCR primers used in this work.

| Primer Number | Sequence | Purpose |
|---|---|---|
| 848 | CCGCCAGGCAAATTCTGTTT | Reverse primer for colony PCR, all sgRNAs |
| 1092 | ACTAGTATTATACCTAGGACTGAGCTAGC | Reverse primer, all sgRNAs |
| 1093 | GTTTTAGAGCTAGAAATAGCAAGTTAAAATAAGGC | Forward primer, control sgRNA |
| 1095 | GCGGGACGACTGAGCTCGCTGTTTTAGAGCTAGAAATAGCAAGTTAAAATAAGGC | Forward primer, sgRNA 9 for targeting *fliF* in K56-2 |
| 1096 | CACGATGCGATTATCGTGACGTTTTAGAGCTAGAAATAGCAAGTTAAAATAAGGC | Forward primer, sgRNA 10 for targeting *fliF* in K56-2 |
| 1097 | CTCCGATCGACCGAAAAGAGGTTTTAGAGCTAGAAATAGCAAGTTAAAATAAGGC | Forward primer, sgRNA 11 for targeting *fliF* in K56-2 |
| 1098 | GTATAATACTAGTGTTTTAGAGCTAGAAATAG | Forward primer for colony PCR, control sgRNA |
| 1100 | TAGGTATAATACTAGTGCGGGAC | Forward primer for colony PCR, sgRNA 9 |
| 1101 | TAGGTATAATACTAGTCACGATGCG | Forward primer for colony PCR, sgRNA 10 |
| 1102 | CCTAGGTATAATACTAGTCTCCGATC | Forward primer for colony PCR, sgRNA 11 |

| | | |
|---|---|---|
| 1161 | GTCGGTTAATAAATTTGTTGGTTTTAGAGCTAG AAATAGCAAGTTAAAATAAGGC | Forward primer, sgRNA 1 for targeting *paaA* in K56-2 |
| 1162 | GTATAAACCAACCGACCGGTGTTTTAGAGCTA GAAATAGCAAGTTAAAATAAGGC | Forward primer, K56-2 sgRNA 2 for targeting *paaA* in K56-2 |
| 1163 | CGAAGTATAAACCAACCGACGTTTTAGAGCTA GAAATAGCAAGTTAAAATAAGGC | Forward primer, K56-2 sgRNA 3 for targeting *paaA* in K56-2 |
| 1164 | TCCTAGGTATAATACTAGTGTCGGTT | Forward primer for colony PCR, sgRNA 1 |
| 1165 | GTGTATAAACCAACCGACCGG | Forward primer for colony PCR, sgRNA 2 |
| 1166 | CGAAGTATAAACCAACCGAC | Forward primer for colony PCR, sgRNA 3 |
| 1171 | CTCCAGATTCTGCCGCAGTGGTTTTAGAGCTAG AAATAGCAAGTTAAAATAAGGC | Forward primer, sgRNA 6 and 15 for targeting *phbC* in K56-2 and ATCC17616, respectively |
| 1172 | CGGGCCGTCATGCTCTGGAGGTTTTAGAGCTAG AAATAGCAAGTTAAAATAAGGC | Forward primer, sgRNA 7 for targeting *phbC* in K56-2 |
| 1173 | CGAGCCGGGCCGTCATGCTCGTTTTAGAGCTAG AAATAGCAAGTTAAAATAAGGC | Forward primer, sgRNA 8 for targeting *phbC* in K56-2 |
| 1174 | CCGTTCAGCCGGGCGTGTTCGTTTTAGAGCTAG AAATAGCAAGTTAAAATAAGGC | Forward primer, sgRNA 21 for targeting *phbC* in E264 |
| 1175 | TTTCCTCGGTGTTGCCTGCGGTTTTAGAGCTAG AAATAGCAAGTTAAAATAAGGC | Forward primer, sgRNA 22 for targeting *phbC* in E264 |

| | | |
|---|---|---|
| 1176 | GTGGTTCCTCGGTGTTTCCTGTTTTAGAGCTAGAAATAGCAAGTTAAAATAAGGC | Forward primer, sgRNA 23 for targeting *phbC* in E264 |
| 1177 | CGGGCCGTCATGCCCTGGAGGTTTTAGAGCTAGAAATAGCAAGTTAAAATAAGGC | Forward primer, sgRNA 16 for targeting *phbC* in ATCC17616 |
| 1178 | CGAGCCGGGCCGTCATGCCCGTTTTAGAGCTAGAAATAGCAAGTTAAAATAAGGC | Forward primer, sgRNA 17 for targeting *phbC* in ATCC17616 |
| 1179 | GTCCTAGGTATAATACTAGTCTCCAG | Forward primer for colony PCR, sgRNA 6 |
| 1180 | CCTAGGTATAATACTAGTCGGGC | Forward primer for colony PCR, sgRNA 7 and 16 |
| 1181 | CCTAGGTATAATACTAGTCGAGCC | Forward primer for colony PCR, sgRNA 8 and 17 |
| 1182 | GGTATAATACTAGTCCGTTCAGCC | Forward primer for colony PCR, sgRNA 21 |
| 1183 | AGTCCTAGGTATAATACTAGTTTTCC | Forward primer for colony PCR, sgRNA 22 |
| 1184 | GGTATAATACTAGTGTGGTTCCTC | Forward primer for colony PCR, sgRNA 23 |
| 1198 | CGACGAATGCCAAGCAGATAGTTTTAGAGCTAGAAATAGCAAGTTAAAATAAGGC | Forward primer, non-target sgRNA in all three species |
| 1199 | CTAGGTATAATACTAGTCGACGAATGC | Forward primer for colony PCR, non-target sgRNA |

| | | |
|---|---|---|
| 1200 | TTAATAAATTTGTTGCGGAATTTGTTTTAGAGCTAGAAATAGCAAGTTAAAATAAGGC | Forward primer, sgRNA 12 for targeting *paaA* in ATCC17616 |
| 1201 | CGCAGTATAAACCAACCGACGTTTTAGAGCTAGAAATAGCAAGTTAAAATAAGGC | Forward primer, sgRNA 13 for targeting *paaA* in ATCC17616 |
| 1202 | ACATGGCGAGCGCTCTCGTCGTTTTAGAGCTAGAAATAGCAAGTTAAAATAAGGC | Forward primer, sgRNA 14 for targeting *paaA* in ATCC17616 |
| 1203 | GTCCTAGGTATAATACTAGTTTAATAAATTTG | Forward primer for colony PCR, sgRNA 12 |
| 1204 | GGTATAATACTAGTCGCAGTATAAAC | Forward primer for colony PCR, sgRNA 13 |
| 1205 | GGTATAATACTAGTACATGGCGAG | Forward primer for colony PCR, sgRNA 14 |
| 1206 | CGGTTAATAAATGATTTTAGGTTTTAGAGCTAGAAATAGCAAGTTAAAATAAGGC | Forward primer, sgRNA 18 for targeting *paaA* in E264 |
| 1207 | GTTCACGCAGTATAAACCAAGTTTTAGAGCTAGAAATAGCAAGTTAAAATAAGGC | Forward primer, E264 sgRNA 19 for targeting *paaA* in E264 |
| 1208 | TGGCGATGTTCTCGTCCAGGGTTTTAGAGCTAGAAATAGCAAGTTAAAATAAGGC | Forward primer, E264 sgRNA 20 for targeting *paaA* in E264 |
| 1209 | GGTATAATACTAGTCGGTTAATAAATGAT | Forward primer for colony PCR, sgRNA 18 |
| 1210 | GGTATAATACTAGTGTTCACGCAG | Forward primer for colony PCR, sgRNA 19 |

| | | |
|---|---|---|
| 1211 | GTATAATACTAGTTGGCGATGTTC | Forward primer for colony PCR, sgRNA 20 |
| 1212 | ACATGGCGAAGGCTCTCGTCGTTTTAGAGCTAGAAATAGCAAGTTAAAATAAGGC | Forward primer, sgRNA 4 for targeting *paaA* in K56-2 |
| 1213 | ACGCAATCCCTCGACATCCCGTTTTAGAGCTAGAAATAGCAAGTTAAAATAAGGC | Forward primer, sgRNA 5 for targeting *paaA* in K56-2 |
| 1214 | GGTATAATACTAGTACATGGCGAAG | Forward primer for colony PCR, sgRNA 4 |
| 1215 | GGTATAATACTAGTACGCAATCCC | Forward primer for colony PCR, sgRNA 5 |

Supplemental Table 4 and 5 have been provided as two separate excel files.

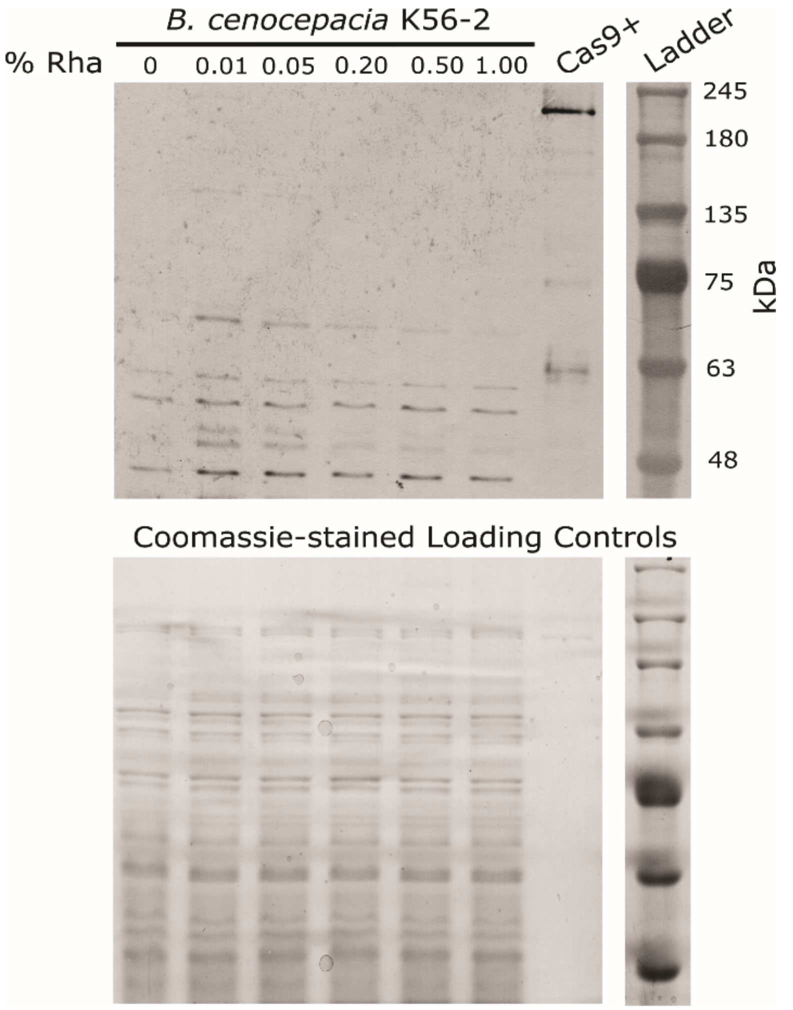


**Supplemental Figure 1. The native *dcas9* is not expressed in *B. cenocepacia* K56-2**. *B. cenocepacia* K56-2::dCas9-native was grown to $OD_{600nm}$ 0.6 then induced fore three hours with the indicated concentrations of rhamnose and the soluble protein fraction was extracted. dCas9 was detected with α-Cas9 antibodies. The Coomassie-stained gel run was in parallel to that used for immunoblotting to ensure equal loading.

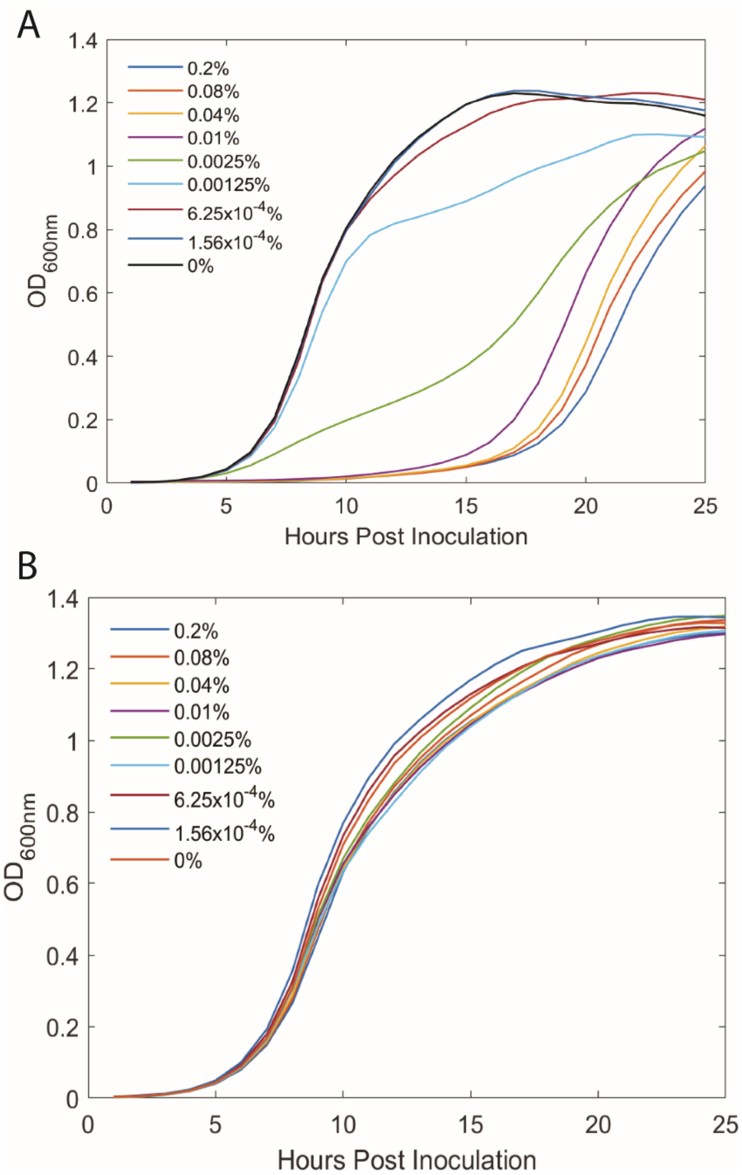


**Supplemental Figure 2. dCas9 expressed from a multicopy plasmid causes a growth defect in *B. cenocepacia* K56-**2. *B. cenocepacia* K56-2 harboring the plasmid pSC-rhadCas9 (A) or the vector control, pSCrhaB2 (B) was cultured with 100 µg/mL trimethoprim and with the indicated concentrations of rhamnose.

A

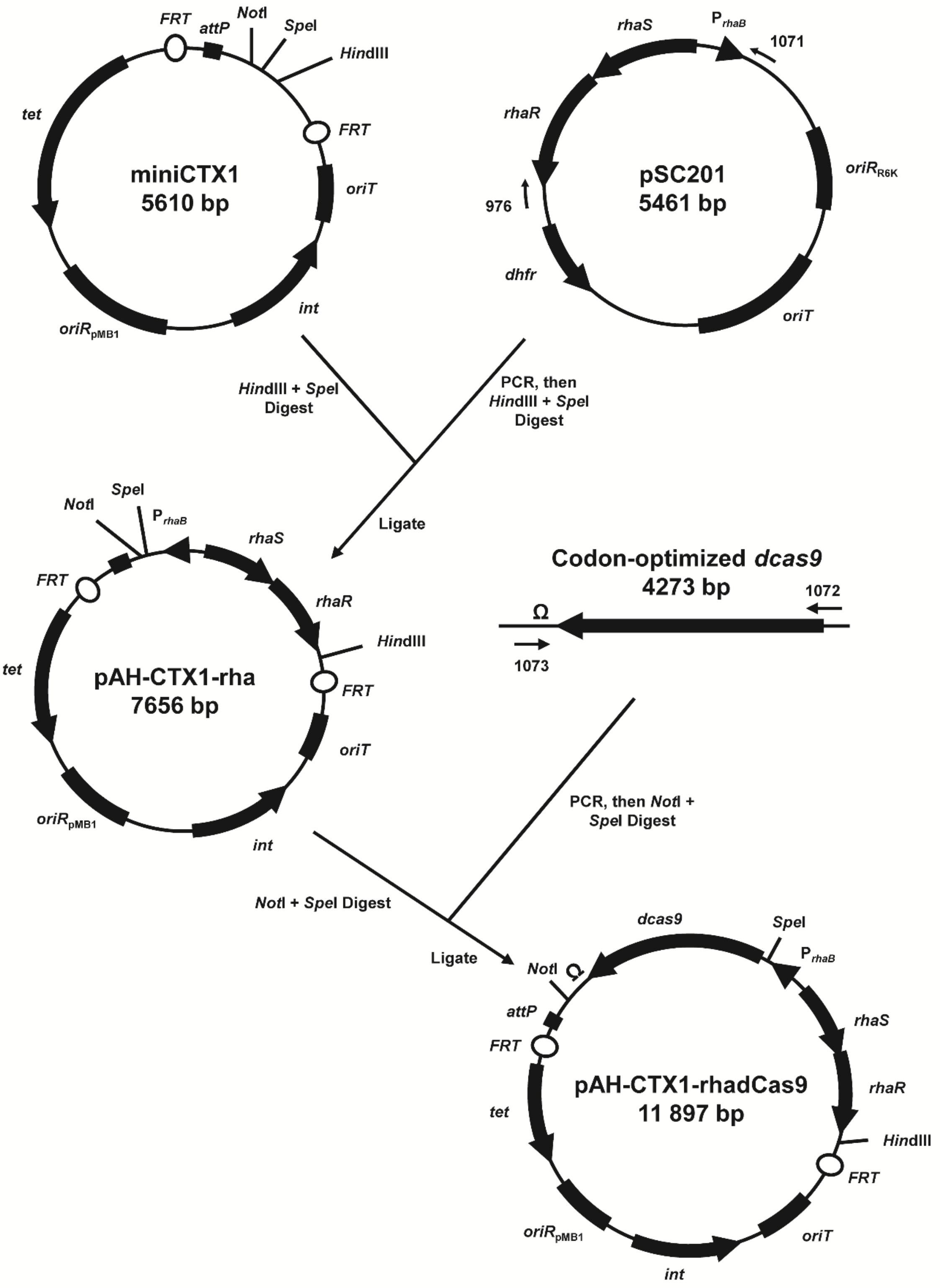

miniCTX1
5610 bp
FRT
attP
NotI
SpeI
HindIII
FRT
oriT
int
oriRpMB1
tet
pSC201
5461 bp
rhaS
PrhaB
1071
oriRR6K
oriT
dhfr
976
rhaR
HindIII + SpeI
Digest
PCR, then
HindIII + SpeI
Digest
Ligate
pAH-CTX1-rha
7656 bp
NotI
SpeI
PrhaB
rhaS
rhaR
HindIII
FRT
oriT
int
oriRpMB1
tet
FRT
Codon-optimized dcas9
4273 bp
Ω
1072
1073
PCR, then NotI +
SpeI Digest
NotI + SpeI Digest
Ligate
pAH-CTX1-rhadCas9
11 897 bp
dcas9
SpeI
PrhaB
rhaS
rhaR
HindIII
FRT
oriT
int
oriRpMB1
tet
FRT
attP
NotI
Ω

B

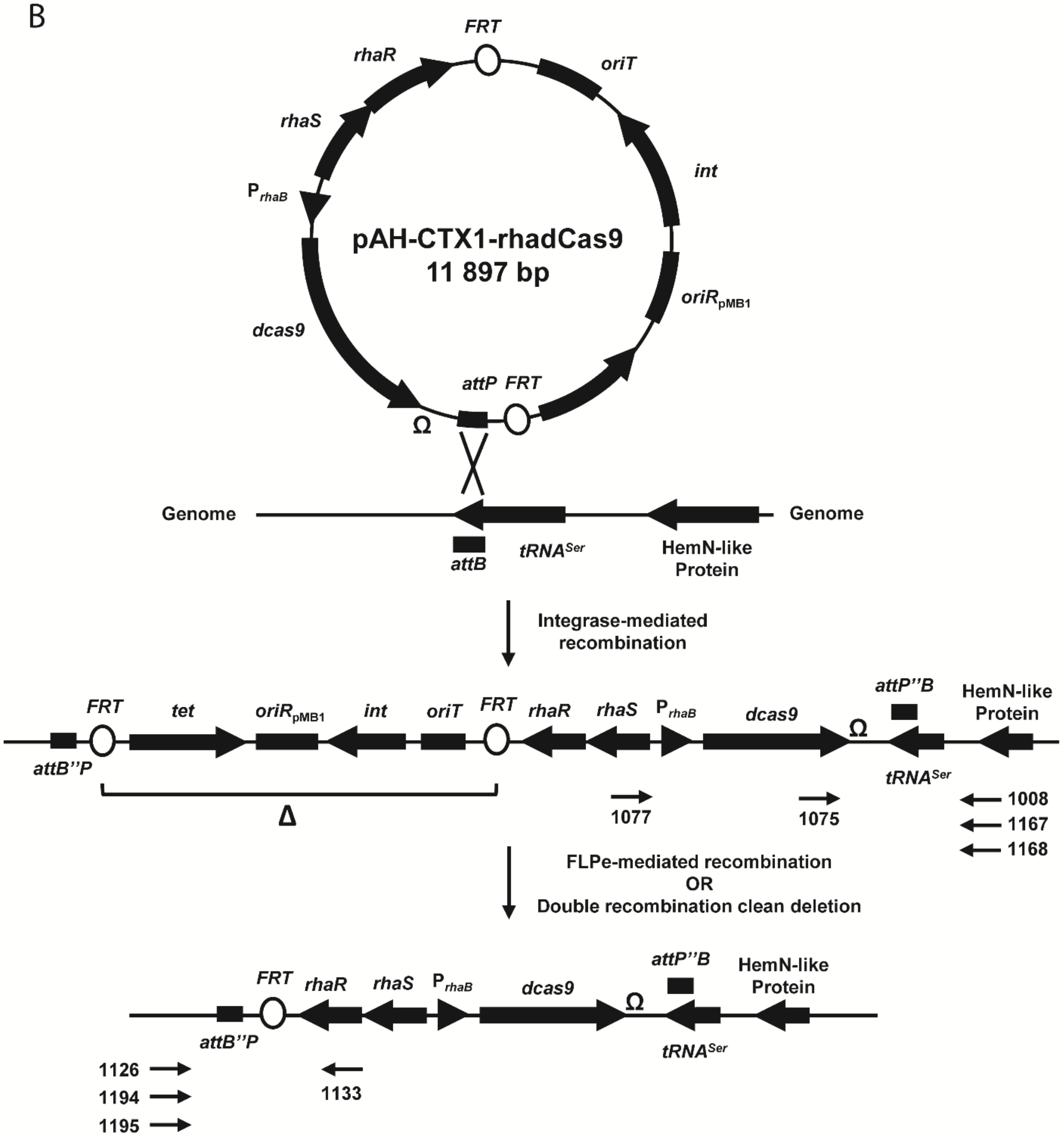


**Supplemental Figure 3. Construction and insertion scheme for the integrative plasmids pAH-CTX1-rha and pAH-CTX1-rhadCas9**. A) The rhamnose-inducible promoter was amplified from pSC201 with *Spe*I and *Hind*III restriction sites and introduced into mini-CTX1, creating pAH-CTX1-rha. The codon-optimized *dcas9* was constructed with flanking *Not*I and *Spe*I restriction sites as per Materials and Methods, and introduced into pAH-CTX1-rha, creating pAH-CTX1-rhadCas9. B) pAH-CTX1-rha and pAH-CTX1-rhadcas9 integrate into the genome of a host organism at the *attB* site, usually located at the 3' end of the serine tRNA. Either FLPe of double homologous recombination can be used to remove the accessory genes, such as *tet*, creating an antibiotic-sensitive clean insertion mutant. Primer numbers used for cloning/colony PCR refer to Supplemental Table 2.

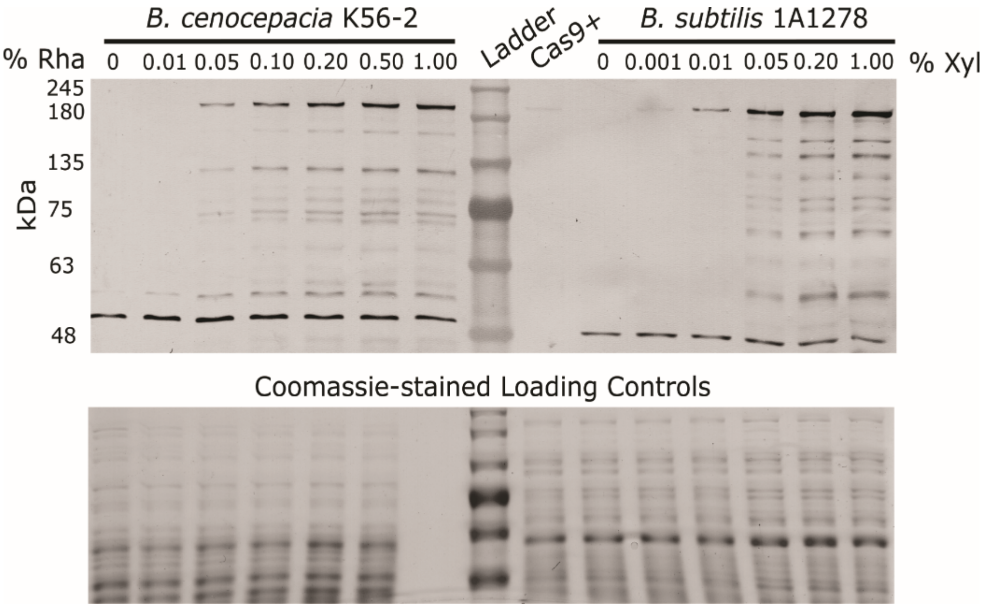


**Supplemental Figure 4. Control experiments for the development of CRISPRi in *Burkholderia*.** Entire immunoblot from Figure 1B and corresponding SDS-PAGE gel stained with Coomassie to demonstrate equal loading.

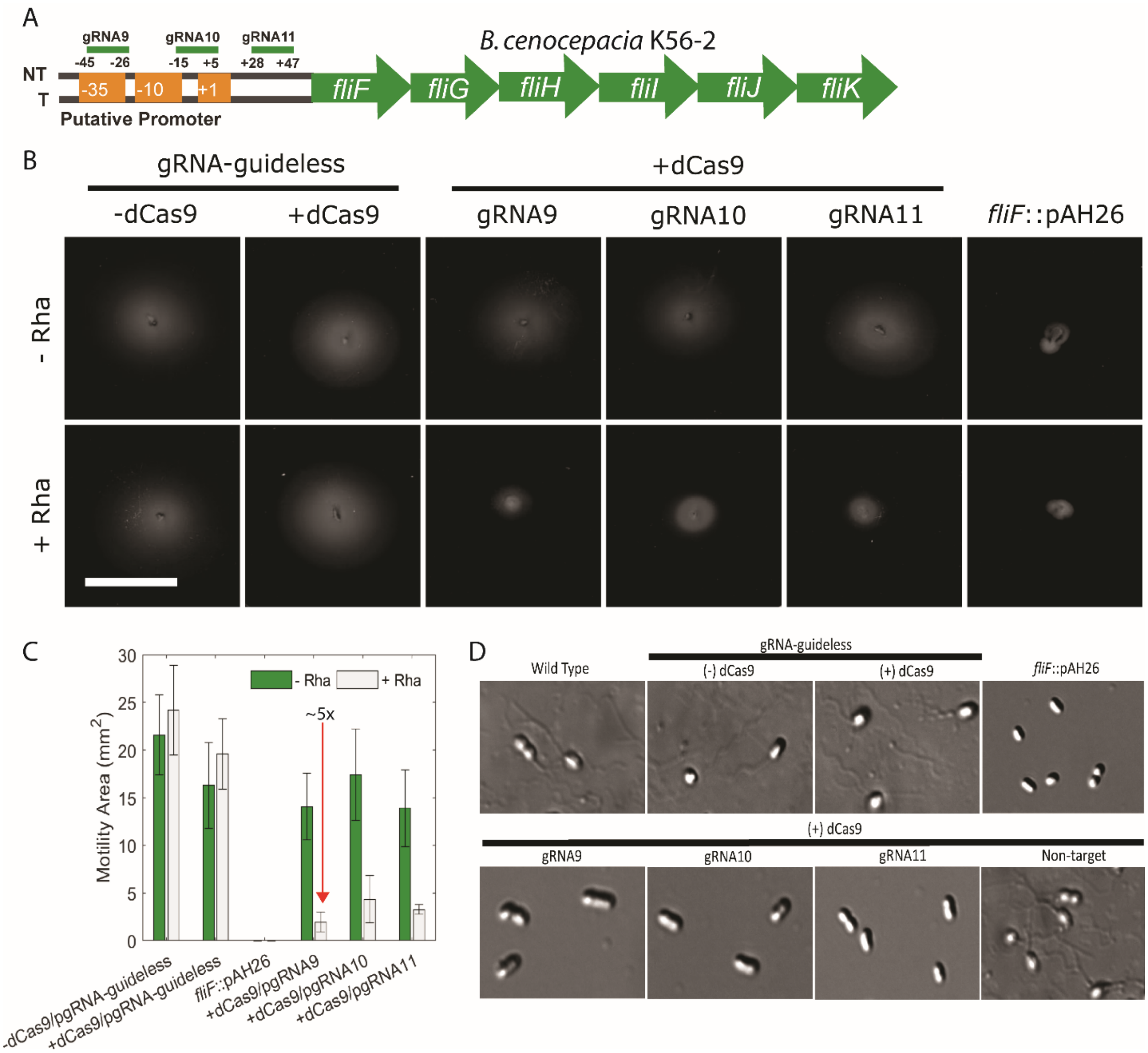


**Supplemental Figure 5. Targeting *fliF* by CRISPRi in *B. cenocepacia* K56-2 reduces swimming motility**. A) Positions of the sgRNAs targeting regions upstream of the *fliFGHIJK* operon. sgRNA9 and 10 were designed to target the promoter elements (-35 and -10 boxes), while sgRNA11 targeted just upstream of the *fliF* ORF on the non-template (NT) strand. B) Motility was reduced, but not ablated by CRISPRi targeting of *fliF*. K56-2::CTX1-rha (-dCas9), K56-2::CTX1-rhadCas9 (+dCas9) (both harbouring pgRNA-guideless), and +dCas9 mutants harbouring sgRNAs targeting *fliF* were stab inoculated into 0.3% agar nutrient broth medium containing 100 µg/mL trimethoprim and with or without 0.2% rhamnose. The scale bar is 2 cm. C) Area of the swimming motility halos in B). D) The indicated strains were grown overnight with 0.2% rhamnose and 100 μg/mL trimethoprim, as required, and used for flagellum staining. Cells were imaged by DIC as we found better contrast versus brightfield microscopy.

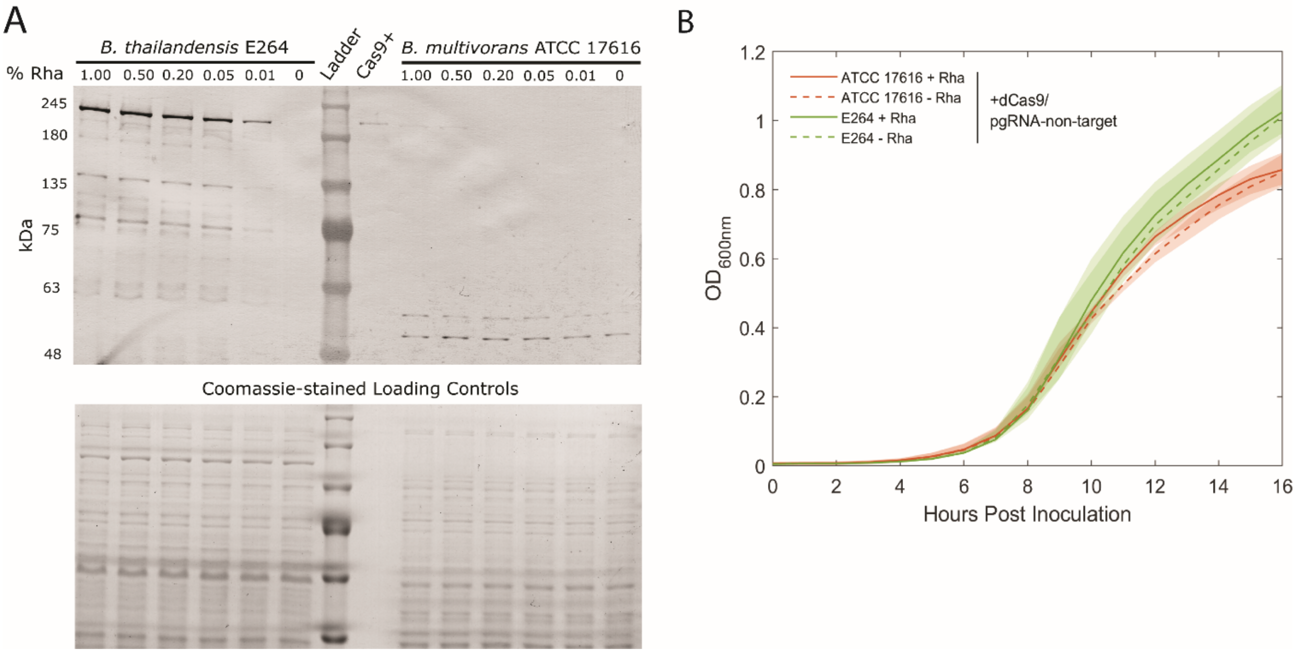


**Supplemental Figure 6. dCas9 expression is detected from chromosomal expression in *B. thailandensis* E264, but poorly from *B. multivorans* ATCC 17616.** A) Strains harbouring the codon-optimized *dcas9* in the chromosome were grown at $OD_{600nm}$ 0.6 then induced for three hours with the indicated concentrations of rhamnose and soluble protein was harvested. dCas9 expression was detected by probing with an α-dCas9 antibody. Below, the protein was run in parallel on SDS-PAGE and stained with Coomassie to ensure equal protein loading. B) Growth curves of *B. multivorans* ATCC17616 and *B. thailandensis* E264 species harbouring a chromosomal copy of the codon-optimized *dcas9* and the non-genome targeting guide plasmid, pgRNA-non-target (Supplemental Table 4).

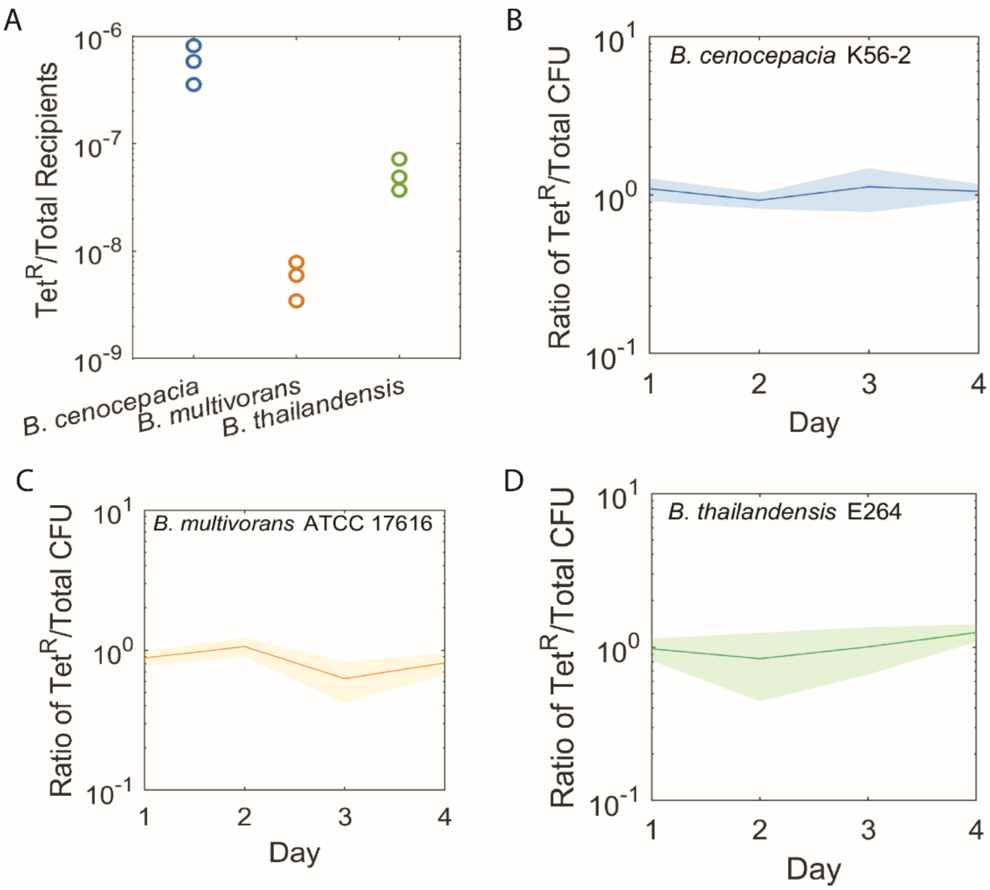


**Supplemental Figure 7. pAH-CTX1-rhadCas9 integrates with variable efficiency but is maintained without antibiotic selection.** A) Strains were subjected to triparental mating with pAH-CTX1-rhadCas9 as per the Materials and Methods. After mating, CFU counts were done on LB agar with 50 µg/mL gentamicin supplemented with and without 50 µg/mL tetracycline (Tet50). Ratios of CFU on the Tet50 to non-selective medium are reported. For B), C), and D) strains harbouring pAH-CTX1-rhadCas9, but not yet cleaned of the plasmid accessory genes, were passaged over four days in LB without tetracycline. For each day, the CFU counts on Tet50 and non-selective medium were determined and the ratios are reported.
